# High-efficiency gene editing in *Anopheles sinensis* using ReMOT control

**DOI:** 10.1101/2023.08.29.555096

**Authors:** Xiao-lin Yang, Xia Ling, Quan Sun, Pin-pin Qiu, Kai Xiang, Jun-feng Hong, Shu-lin He, Jie Chen, Xin Ding, Hai Hu, Zheng-bo He, Cao Zhou, Bin Chen, Liang Qiao

## Abstract

CRISPR/Cas9-mediated gene editing provides an effective method for deciphering the molecular mechanisms underlying mosquito development and mosquito-borne disease transmission, as well as for exploring genetic control strategies. However, delivering the Cas9 ribonucleoprotein complex by embryo injection to produce genetic modifications is challenging, is mostly confined to model mosquitoes and specialized laboratories, and has low editing efficiency. Here, we established an effective Receptor-Mediated Ovary Transduction of Cargo (ReMOT) control method, enabling the introduction of heritable mutations into *Anopheles sinensis*, the major malaria vector in China and Southeast Asia, via the injection of female adult mosquitoes. Injection of a mixture of P2C-DsRed and saponin resulted in red fluorescence in the ovaries, with a 100% success rate. Using this system, we knocked-out the pigment synthesis genes, *Aswhite* and *Asyellow*, using injected wild-type (WT) females mated with WT males, resulting in the highest efficiency of gene editing among mosquitoes under the same mating conditions. Furthermore, the gene-editing efficiency was increased by at least 2.1-fold using injected WT females mated with mutant males. This improved ReMOT control method exhibits high editing efficiency, with important benefits in terms of functional genomics research and genetic control strategies in *An. sinensis*. Moreover, this represents a convenient method for gene manipulation in laboratories that are unable to perform embryo injection or that lack embryo-injection equipment.

## Introduction

Functional genomics studies of mosquito vectors are crucial for understanding their physiological behavior, pathogen transmission, and evolution, and for providing essential molecular targets for genetic control (Severson & Behura, 2011; Ruzzante *et al*., 2019; Alphey, 2014; HONG *et al*., 2022). The main methods involve CRISPR-Cas9 gene editing via embryo microinjection (2016; Criscione *et al*., 2015; Kistler *et al*., 2015); however, technical barriers, including oviposition types, injection timing, training, expensive equipment, and specific injection protocols for different mosquito species (Criscione *et al*., 2015; Au-Carballar-Lejarazú *et al*., 2021), represent difficulties for laboratories studying non-model mosquitoes or lacking the required expertise and facilities for microinjection. The Receptor-Mediated Ovary Transduction of Cargo (ReMOT) method has been employed for germline-level gene editing in some model mosquitoes, involving injection of a Cas9/gRNA complex fused with a P2C ovarian-targeting peptide into adult female mosquitoes (Chaverra-Rodriguez *et al*., 2018; Macias *et al*., 2020; Li *et al*., 2021). Its convenience and safety mean that ReMOT provides an alternative to embryonic injection; however, its general applicability and efficiency remain unclear.

*Anopheles sinensis* is an important malaria vector in China and Southeast Asia (Feng *et al*., 2017; Zhang *et al*., 2017). Although its genome has been reported (Zhou *et al*., 2014), progress in suitable genomics techniques has been slow. Furthermore, the inability of embryo injection to achieve high-efficiency editing and its technical challenges make it unsuitable for large-scale and efficient gene-function surveys (Liu *et al*., 2021), thus limiting studies to improve our understanding of the physiological development of this species and impeding the development of genetic control strategies.

## Results and discussion

This study revealed that *AsVg* transcription increased significantly from 12 to 20 h post-blood-feeding (Fig. 1A), indicating high levels of vitellogenesis at this stage (Macias *et al*., 2020; Sappington *et al*., 1995). In addition, *A. sinensis* eggs undergo rapid growth from 20 to 40 h post-blood-feeding, reaching a relatively mature state (Fig. 1B). This stage signifies an absorption peak of ovarian cells via receptor-mediated endocytosis (Sappington *et al*., 1995; Izumi *et al*., 1994). Studies in other species found that the highest editing efficiency was achieved by injecting P2C-ribonucleoprotein (RNP) complexes during vitellogenesis (Chaverra-Rodriguez *et al*., 2018; Macias *et al*., 2020; Li *et al*., 2021; Shirai *et al*., 2022). We therefore hypothesized that injection at 20 h post-blood-feeding might improve the transportation and accumulation of P2C-RNP complexes in the eggs, resulting in higher editing efficiency.

**Figure 1.**
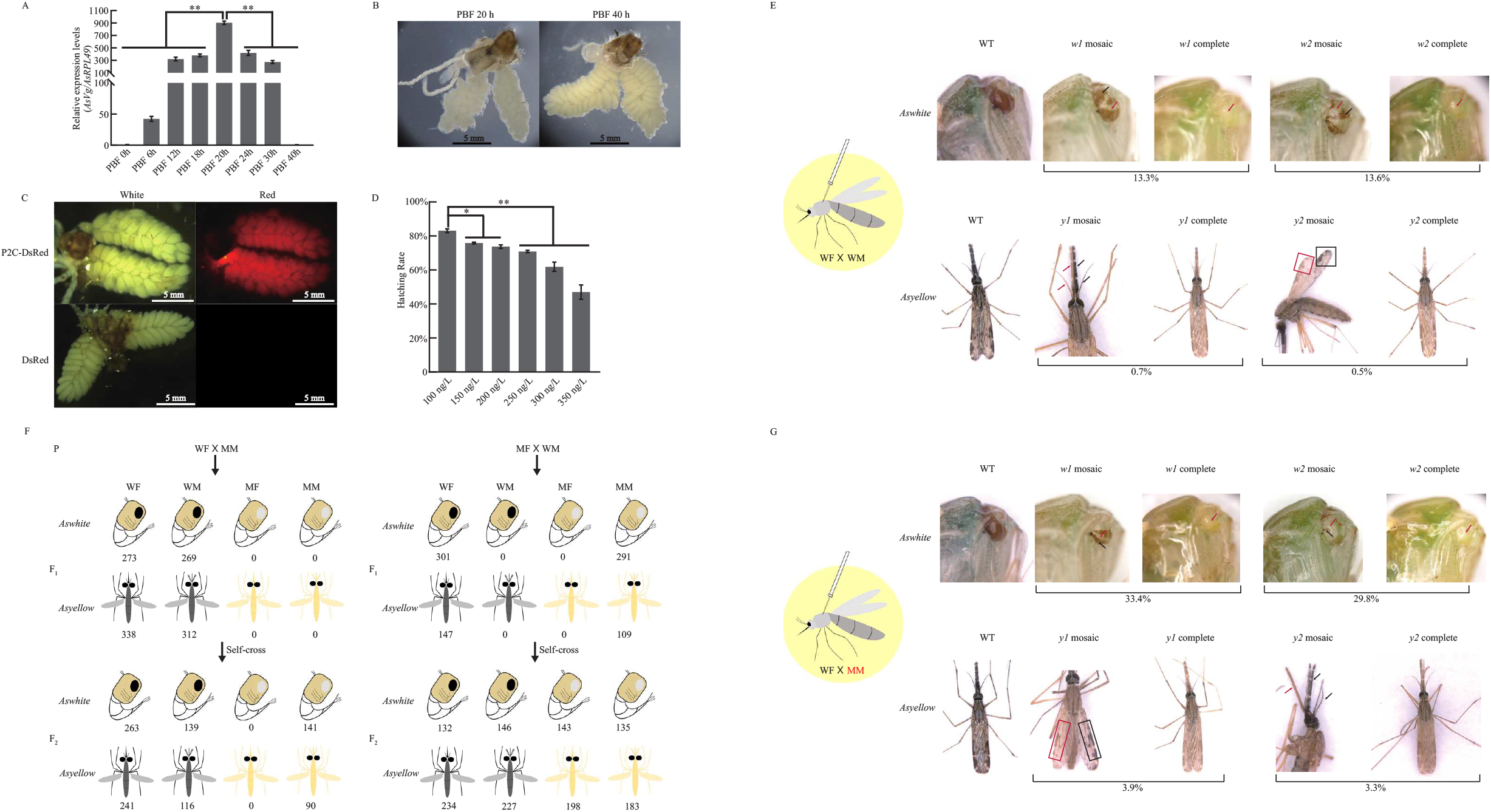
ReMOT control gene-editing system in *Anopheles sinensis*. A. Expression patterns of *AsVg* transcripts at different developmental stages post-blood-feeding (PBF) in female mosquitoes. *AsRPL49* used as an internal control. Statistical analysis performed using *t*-tests (n = 3); \*\**p*<0.01. B. Morphology of ovarian tissues at 20 and 40 h PBF in female mosquitoes. Scale bar = 5 mm. C. Fluorescent phenotypes of ovarian tissues after injection of P2C-DsRed (treatment group) into female mosquitoes. DsRed recombinant proteins used as control. Scale bar = 5 mm. D. Hatching rates of offspring of mosquitoes injected with saponin plus P2C-Linker-Cas9 recombinant protein at different concentrations. Statistical analysis performed using *t*-tests (n = 2); \**p*<0.05, \*\**p*<0.01. E. Phenotypic characteristics of offspring of WT females injected with RNP complexes mated with WT males. Red arrows and boxes, mutated characteristics; black arrows and boxes, WT characteristics at corresponding sites in mosaic individuals. F. Sex-linked inheritance of *Aswhite* and *Asyellow* genes. WF, WT female; MM, mutant male; WM, WT male; MF, mutant female. Under the WF×MM mating scheme, the sex ratio and phenotype distribution in the F_2_ generation indicated that the mutant gene did not show conventional autosomal genetic inheritance (*Aswhite*: χ2 = 184.48, df =3 \*\**p*<0.01; *Asyellow*: χ ^2^= 124.63, df = 3, \*\**p*<0.01). Similarly, under the MF×WM mating scheme, the sex ratio and phenotype distribution in F_1_ generations deviated significantly from the pattern observed under the WF×MM mating scheme, and inheritance of the mutant gene did not show conventional autosomal genetic inheritance (*Aswhite*: χ^2^ = =587.08, df = 3, \*\**p*<0.01; *Asyellow*: χ^2^= 239.82, df = 3, \*\**p*<0.01). G. Phenotypic characteristics of offspring of WF mated with MM. Red arrows and boxes, mutated phenotypic characteristics; black arrows and boxes, WT phenotypic characteristics at corresponding sites in mosaic individuals.

Injection of P2C-DsRed protein (Fig. S1) resulted in red fluorescence in the ovaries in all individuals, compared with the control group (injection of DsRed protein resulted in none fluorescence) (Fig. 1C, Table S2). Ovarian tissue sections and western blot analyses confirmed significant enrichment of red fluorescent protein in the eggs in the P2C-DsRed-injection group (Fig. S2). These results indicated that P2C facilitated the uptake of exogenous proteins into the ovaries in *A. sinensis*. Furthermore, the hatching rate following injection of saponin plus P2C-Linker-Cas9 protein (Fig. S1) was highest at a saponin concentration of 100 ng/μL (Fig. 1D, Table S3), resulting in ample progeny for phenotypic observations.

We used *kynurenine 3-monooxygenase* (‘*white*’) (Han *et al*., 2003) and *dopachrome conversion enzyme* (‘*yellow*’) (Wittkopp *et al*., 2002) as marker genes to determine the efficiency of ReMOT editing in *An. sinensis. In vitro* cleavage assays showed cleavage of target sites by the RNP complexes in both the treatment and control groups (Figs. S1, S3). Mating wild-type (WT) males with WT females resulted in mean gene-editing efficiencies for the *w1* and *w2* target sites of *Aswhite* of 13.3% and 13.6%, respectively (Fig. 1E, Table S4, Supplemental file 1-2). This represented an increase of ≥33-fold in editing efficiency compared with the outcomes in other model mosquitoes employing the same mating approach (Table S4). Furthermore, self-crossing of G_0_ generation mosaic and white-eyed mosquitoes both yielded G_1_ generation white-eyed mutant phenotypes (Table S5), indicating germline-level gene editing.

Mutant phenotypes also occurred in the control group, but with an incidence <1.2%. In addition, these mutations did not appear in the offspring after parental self-crossing, suggesting that they originated from somatic-cell editing (Table S5). The delivery of Cas9-gRNA complexes without P2C (referred to as DIPA-Cas9) achieved successful gene editing in *Blattella germanica* and *Aedes aegypti* (Shirai *et al*., 2022; Shirai *et al*., 2023) but low yields in *Anopheles* (Macias *et al*., 2020), showing that gene editing involving DIPA-Cas9 is unsuitable for use in *Anopheles*, and underscoring the importance of ovary-targeting peptides for the directed delivery of RNPs in these mosquitoes.

Following *Asyellow* knockout (KO), the phenotypes of *y1*-KO and *y2*-KO individuals (injected WT females mated with WT males) included overall yellow coloration, as well as localized mosaic yellow coloring in areas such as the mouthparts, antennae and wings (Fig. 1E), with gene-editing efficiency values of 0.7% and 0.5%, respectively (Table S6, Supplemental file 1-2). No G_0_ generation mutant-phenotype individuals occurred in the control group. In addition, the phenotypes of mutant individuals in the treatment group were heritable (Table S5).

Outcross experiments indicated that mating mutant males with WT females, as well as mutant females with WT males, resulted in a discrepancy between phenotypic sex and phenotype ratio in the F_1_ and F_2_ generations (Fig. 1F), conforming to the principle of sex-linked inheritance and confirming that both *Aswhite* and *Asyellow* were located on the X chromosome. The offspring of WT female and WT male matings could thus potentially be heterozygous for edited and WT alleles, thereby masking the mutant phenotype, potentially leading to underestimation of the actual editing efficiency.

Injections should thus be performed in WT females mated with mutant males. Once the target-gene locus in the female ovary cells is edited, the offspring can exhibit the corresponding mutant phenotype, avoiding the masking effect of heterozygotes and further correcting the editing efficiency. This increased the mean editing efficiencies of the *w1* and *w2* sites of *Aswhite* to 33.4% and 29.8% (Fig. 1G, Table S7, Supplemental file 1-2), respectively, which were approximately 5.3-fold higher than the editing efficiency in other model mosquitoes under the same mating conditions (Table S7). The editing efficiencies at the *y1* and *y2* sites of *Asyellow* were 3.9% and 3.3% (Fig. 1G, Table S8, Supplemental file 1-2), respectively, which were only lower than that observed in *Anopheles stephensi* under certain injection conditions (Table S8). Compared with the injection-treated group, a few mutants were observed in the control group, but no heritable mutant types occurred in their self-crossed offspring (Table S9).

The hatching and eclosion rates of *Asyellow* mutants were significantly lower compared with WT mosquitoes (data not shown). Similar results were previously noted in *Aedes albopictus* and *Agrotis ypsilon* (Noh *et al*., 2020; Noh *et al*., 2021; Chen *et al*., 2018). In addition, mosaic phenotypes in small sclerotized regions are difficult to observe. Moreover, functional redundancy among members of the *yellow* gene family might lead to compensatory effects for mutant phenotypes (Noh *et al*., 2020; Li & Christensen, 2011). These factors may collectively reduce the editing efficiency for *Asyellow* compared with *Aswhite*.

## Conclusion

In summary, we established a high-efficiency gene-editing method in *A. sinensis* using a ReMOT control strategy. Our results validated the applicability and convenience of ReMOT for functional genomics research in non-model mosquitoes, as well as providing valuable insights for functional genomics techniques in other insects, and for laboratories lacking the capability for mosquito embryo microinjection.

## Acknowledgments

This work was supported by the National Natural Science Foundation of China (Nos. 31772527; 31872262), the Natural Science Foundation Project of Chongqing (No. CSTB2022NSCQ-MSX1355), the Scientific and Technological Research Program of Chongqing Municipal Education Commission (KJZD-K202200507, KJQN202200533); the BAYU scholar program (YS2019027), the Venture and Innovation Support Program for Chongqing Overseas Returnees (No. cx2022052) and the Graduate Research and Innovation Foundation of Chongqing under Grant (CYS23399).

## Figure legends

Fig. S1. Prokaryotic expression and immunoblotting detection of recombinant proteins. Red box indicates target recombinant protein in supernatant lysate of *Escherichia coli*; red arrow indicates western blotting result of target recombinant protein. Lane A: lysate of *E. coli*; Lane B: flow-through lysate after binding to Ni-NTA agarose.

Fig. S2. Fluorescence analysis of ovarian tissue slices from adult mosquitoes injected with P2C-DsRed and DsRed (A), and immunoblot analysis of recombinant protein enrichment in ovarian tissue (B).

Fig. S3. *In vitro* cleavage assays. *In vitro* cleavage of (A) P2C-Linker-Cas9-gRNA RNP complexes, and (B) pET28a-Cas9-gRNA complexes. C. Control cleavage reaction with recombinant protein without gRNA. T, treatment group containing RNP complexes for cleavage system. White arrows indicate intact template DNA; red arrows indicate DNA fragments resulting from cleavage by RNP complexes.

