## Supplementary material for "High-efficiency gene editing in *Anopheles sinensis* using ReMOT control": 20230821 Supplemental file 1 types of mutant sequences-1.pdf

### *w1* site

| | | | |
|---|---|---|---|
| WT | | TCAACACCGATGACCAGTACGGGGATGGAGAGAAT | |
| Mutant-1 | +1 | TCAACACCGATGCACCAGTACGGGGATGGAGAGAAT | $G_0$ from WT females mated with WT males |
| Mutant-2 | -20 | TCAACACCGATA--------------------AAT |  |
| Mutant-3 | +4 | TCAACACCGATGTCATACCAGTACGGGGATGGAGAGAAT | |
| Mutant-4 | -1/+2 | TCAACACCGAT--TAACCGGTACGGGGATGGAGAGAAT | |
| Mutant-5 | -3/+23 | TCAACACCG---GGGCTGTGTTGTAAAATAGATAAACCAGTACGGGGATGGAGAGAAT | |

| | | | |
|---|---|---|---|
| Mutant-1 | -1/+15 | TCAACACCGAT-TCGAACCGTTGGAACACCAGTACGGGGATGGAGAGAAT | $G_0$ from WT females mated with mutant males |
| Mutant-2 | -5/+13 | TCAACACCGAT-----TGGAATCAACACCGTACGGGGATGGAGAGAAT |  |
| Mutant-3 | -7 | TCAACACC-------AGTACGGGGATGGAGAGAAT |  |
| Mutant-4 | -2 | TCAACACCGA--ACCAGTACGGGGATGGAGAGAAT | |
| Mutant-5 | -10 | TCAACA----------TGTACGGG--ATGGAGAGAAT |  |

### *w2* site

| | | | |
|---|---|---|---|
| WT | | CACGAGCGGCTCACGTACACCTGGAAGGAGAT | |
| Mutant-1 | -8 | CACGAGCGGCTCAC--------GGAAGGAGAT | $G_0$ from WT females mated with WT males |
| Mutant-2 | -4 | CACGAGCGGCTCACGTAA----GGAAGGAGAT |  |
| Mutant-3 | +3 | CACGAGCGGCTCACGTACACCGTACGGAAGGAG | |
| Mutant-4 | -6 | CACGAGCGGCTCACGTAC------AAGGAGAT |  |
| Mutant-5 | -2 | CACGAGCGGCTCACGTACAC--GGAAGGAGAT | |

| | | | |
|---|---|---|---|
| Mutant-1 | -3 | CACGAGCGGCTCACGTACATC---AAGGAGAT | $G_0$ from WT females mated with mutant males |
| Mutant-2 | -3 | CACGAGCGGCTCACGTACAC---GAAGGAGAT | |
| Mutant-3 | -1/+2 | CACGAGCGGCTCACGTACACCT-TCGAAGGAGAT | |
| Mutant-4 | -5 | CACGAGCGGCTCACG-----GCGGAAGGAGAT |  |
| Mutant-5 | -1/+3 | CACGAGCGGCTCACGTACACC-ACGGGAAGGAG | |

Blue letters indicate the PAM sequence; Red letters indicate the sgRNA template sequence; "-" symbol indicates deletion of base; Green letter indicates insertion of base; brown letter indicates substitution of base

### *y1* site

| | | |
|---|---|---|
| WT | | GCTAGCGGGGACTACGTCCCGACCAACGGTCTGCCGGTCGGCATCGAGCGCT |
| Mutant-1 | -5/+15 | GCTAGCGGGGACTACGTCCCGACCAAC-----CGGCTACGACCGAAGGCCGGTCGGCATC |
| Mutant-2 | +2 | GCTAGCGGGGACTACGTCCCGACCAACGGTCGATGCCGGTCGGCATCGAGCGCT |
| Mutant-3 | -1/+5 | GCTAGCGGGGACTACGTCCCGACCAACGGTC-GAACGGCCGGTCGGCATCGAGCGCT |
| Mutant-4 | -8 | GCTAGCGGGGACTACGTCCCGACCAACGG--------TCGGCATCGAGCGCT |

$G_0$ from WT females mated with WT males

| | | |
|---|---|---|
| Mutant-1 | -8 | GCTAGCGGGGACTACGTCCCGACCAACGC--------TCGGCATCGAGCGCT |
| Mutant-2 | -14 | GCTAGCGGGGACTACGTTCCGACCAACGGTC--------------GAACGCT |
| Mutant-3 | -4 | GCTAGCGGGGACTACGTCCCGACCAACCA----CCGGTCGGCATCGAGCGCT |
| Mutant-4 | -8 | GCTAGCGGGGACTACGTCCCGACCAACGGTCGAA--------ATCGAGCGCT |
| Mutant-5 | - 4 | GCTAGCGGGGACTACGTCCCGACCAACGGTC----GGTCGGCATCGAGCGCT |

$G_0$ from WT females mated with mutant males

### *y2* site

| | | |
|---|---|---|
| WT | | GCGTACATGTCCGACGAACTGGGCTACGGTCTGATCGTGT |
| Mutant-1 | -3/+15 | GCGTACATGTCCGACGAACTGG---CAGAAATATCAACAAACGGTCTG |
| Mutant-2 | -5 | GCGTACATGTCCGACGAACG-----ACGGTCTGATCGTGT |
| Mutant-3 | -5 | GCGTACATGTCCGACGAACT-----ACGGTCTGATCGTGT |
| Mutant-4 | -7 | GCGTACATGTCCGACGAACT-------GGTCTGATCGTGT |
| Mutant-5 | +1 | GCGTACATGTCCGACGAACTGGGTCTACGGTCTGATCGTGT |

$G_0$ from WT females mated with WT males

| | | |
|---|---|---|
| Mutant-1 | -1/+6 | GCGTACATGTCCGACGAACTGG-TGGGCACGACGGTCTGATCGTGT |
| Mutant-2 | -11 | GCGTACATGTCCGACGAACTGGG-----------TCGTGT |
| Mutant-3 | - 5 | GCGTACATGTCCGACGAACTGG-----GGTCTGATCGTGT |
| Mutant-4 | -19 | GCGTACATGTCCGACGAA-------------------TGT |
| Mutant-5 | -2 | GCGTACATGTCCGACGGTACGTAC--CGGTCTGATCGTGT |

$G_0$ from WT females mated with mutant males

Blue letters indicate the PAM sequence; Red letters indicate the sgRNA template sequence; “-” symbol indicates deletion of base; Green letter indicates insertion of base; brown letter indicates substitution of base
