## Supplementary material for "High-efficiency gene editing in *Anopheles sinensis* using ReMOT control": 20230821 Supplemental file 2 sequence information.pdf

>Aswhite-sgRNA1-WT
ACGGCAGCGGTTGGAGACGCCCAGTGCAGAGCTGCTGGGGAAAAGTGCAGAAGCAA
ATAACCGATTTCTGCGGTTCATAACCGAAGCGCTACAAGGTTAAAAAGTGACTTGTAGC
TCCGTTTGCGAACGGGTCAGTCATCGAGGACCTCATATCTTACGTTAAAGACAGCCGG
AATCAAGCAGTAATAATACCATGACAATCAACACCGATGACCAGTACGGGGATGGAGA
GAATAAATCCACTATCAGCTCCAGTCGGGTAAGTCTGAGGTTCACAAGCGTTCTCGTTT
TGATAGCAGCAGGTAAAAATGCTGGAAAATGTATTGGGTGCTACTGTTTCCCGAGAAG
TATCACATCACATCATATAGCAGATCTACTATTGTTTGGAGTTAAGGAGAGTAGATTTAA
AAGGCA
>Aswhite-sgRNA1-mutation 1 (WT females × WT males)
ACGGCAGCGGTTGGAGACGCCCAGTGCAGAGCTGCTGGGGAAAAGTGCAGAAGCAA
ATAACCGATTTCTGCGGTTCATAACCGAAGCGCTACAAGGTTAAAAAGTGACTTGTAGC
TCCGTTTGCGAACGGGTCAGTCATCGAGGACCTCATATCTTACGTTAAAGACAGCCGG
AATCAAGCAGTAATAATACCATGACAATCAACACCGATGCACCAGTACGGGGATGGAG
AGAATAAATCCACTATCAGCTCCAGTCGGGTAAGTCTGAGGTTCACAAGCGTTCTCGTT
TTGATAGCAGCAGGTAAAAATGCTGGAAAATGTATTGGGTGCTACTGTTTCCCGAGAA
GTATCACATCACATCATATAGCAGATCTACTATTGTTTGGAGTTAAGGAGAGTAGATTTA
AAAGGCA
>Aswhite-sgRNA1-mutation 2 (WT females × WT males)
ACGGCAGCGGTTGGAGACGCCCAGTGCAGAGCTGCTGGGGAAAAGTGCAGAAGCAA
ATAACCGATTTCTGCGGTTCATAACCGAAGCGCTACAAGGTTAAAAAGTGACTTGTAGC
TCCGTTTGCGAACGGGTCAGTCATCGAGGACCTCATATCTTACGTTAAAGACAGCCGG
AATCAAGCAGTAATAATACCATGACAATCAACACCGATAAATCAATCCACTATCAGCTC
CAGTCGGGTAAGTCTGAGGTTCACAAGCGTTCTCGTTTTGATAGCAGCAGGTAAAAAT
GCTGGAAAATGTATTGGGTGCTACTGTTTCCCGAGAAGTATCACATCACATCATATAGC
AGATCTACTATTGTTTGGAGTTAAGGAGAGTAGATTTAAAAGGCA
>Aswhite-sgRNA1-mutation 3 (WT females × WT males)
ACGGCAGCGGTTGGAGACGCCCAGTGCAGAGCTGCTGGGGAAAAGTGCAGAAGCAA
ATAACCGATTTCTGCGGTTCATAACCGAAGCGCTACAAGGTTAAAAAGTGACTTGTAGC
TCCGTTTGCGAACGGGTCAGTCATCGAGGACCTCATATCTTACGTTAAAGACAGCCGG
AATCAAGCAGTAATAATACCATGACAATCAACACCGATGTCATACCAGTACGGGGATGG
AGAGAATAAATCCACTATCAGCTCCAGTCGGGTAAGTCTGAGGTTCACAAGCGTTCTC
GTTTTGATAGCAGCAGGTAAAAATGCTGGAAAATGTATTGGGTGCTACTGTTTCCCGAG
AAGTATCACATCACATCATATAGCAGATCTACTATTGTTTGGAGTTAAGGAGAGTAGATT
TAAAAGGCA
>Aswhite-sgRNA1-mutation 4 (WT females × WT males)
ACGGCAGCGGTTGGAGACGCCCAGTGCAGAGCTGCTGGGGAAAAGTGCAGAAGCAA
ATAACCGATTTCTGCGGTTCATAACCGAAGCGCTACAAGGTTAAAAAGTGACTTGTAGC
TCCGTTTGCGAACGGGTCAGTCATCGAGGACCTCATATCTTACGTTAAAGACAGCCGG
AATCAAGCAGTAATAATACCATGACAATCAACACCGATTAACCGGTACGGGGATGGAG
AGAATAAATCCACTATCAGCTCCAGTCGGGTAAGTCTGAGGTTCACAAGCGTTCTCGTT
TTGATAGCAGCAGGTAAAAATGCTGGAAAATGTATTGGGTGCTACTGTTTCCCGAGAA
GTATCACATCACATCATATAGCAGATCTACTATTGTTTGGAGTTAAGGAGAGTAGATTTA
AAAGGCA

>Aswhite-sgRNA1-mutation 5 (WT females × WT males)
ACGGCAGCGGTTGGAGACGCCCAGTGCAGAGCTGCTGGGGAAAAGTGCAGAAGCAA
ATAACCGATTTCTGCGGTTCATAACCGAAGCGCTACAAGGTTAAAAAGTGACTTGTAGC
TCCGTTTGCGAACGGGTCAGTCATCGAGGACCTCATATCTTACGTTAAAGACAGCCGG
AATCAAGCAGTAATAATACCATGACAATCAACACCGGGGCTGTGTTGTAAAATAGATAA
ACCAGTACGGGGATGGAGAGAATAAATCCACTATCAGCTCCAGTCGGGTAAGTCTGAG
GTTCACAAGCGTTCTCGTTTTGATAGCAGCAGGTAAAAATGCTGGAAAATGTATTGGGT
GCTACTGTTTCCCGAGAAGTATCACATCACATCATATAGCAGATCTACTATTGTTTGGAG
TTAAGGAGAGTAGATTTAAAAGGCA
>Aswhite-sgRNA1-mutation 1 (WT females × mutant males)
ACGGCAGCGGTTGGAGACGCCCAGTGCAGAGCTGCTGGGGAAAAGTGCAGAAGCAA
ATAACCGATTTCTGCGGTTCATAACCGAAGCGCTACAAGGTTAAAAAGTGACTTGTAGC
TCCGTTTGCGAACGGGTCAGTCATCGAGGACCTCATATCTTACGTTAAAGACAGCCGG
AATCAAGCAGTAATAATACCATGACAATCAACACCGATTCGAACCGTTGGAACACCAG
TACGGGGATGGAGAGAATAAATCCACTATCAGCTCCAGTCGGGTAAGTCTGAGGTTCA
CAAGCGTTCTCGTTTTGATAGCAGCAGGTAAAAATGCTGGAAAATGTATTGGGTGCTAC
TGTTTCCCGAGAAGTATCACATCACATCATATAGCAGATCTACTATTGTTTGGAGTTAAG
GAGAGTAGATTTAAAAGGCA
>Aswhite-sgRNA1-mutation 2 (WT females × mutant males)
ACGGCAGCGGTTGGAGACGCCCAGTGCAGAGCTGCTGGGGAAAAGTGCAGAAGCAA
ATAACCGATTTCTGCGGTTCATAACCGAAGCGCTACAAGGTTAAAAAGTGACTTGTAGC
TCCGTTTGCGAACGGGTCAGTCATCGAGGACCTCATATCTTACGTTAAAGACAGCCGG
AATCAAGCAGTAATAATACCATGACAATCAACACCGATTGGAATCAACACCGTACGGG
GATGGAGAGAATAAATCCACTATCAGCTCCAGTCGGGTAAGTCTGAGGTTCACAAGCG
TTCTCGTTTTGATAGCAGCAGGTAAAAATGCTGGAAAATGTATTGGGTGCTACTGTTTC
CCGAGAAGTATCACATCACATCATATAGCAGATCTACTATTGTTTGGAGTTAAGGAGAG
TAGATTTAAAAGGCA
>Aswhite-sgRNA1-mutation 3 (WT females × mutant males)
ACGGCAGCGGTTGGAGACGCCCAGTGCAGAGCTGCTGGGGAAAAGTGCAGAAGCAA
ATAACCGATTTCTGCGGTTCATAACCGAAGCGCTACAAGGTTAAAAAGTGACTTGTAGC
TCCGTTTGCGAACGGGTCAGTCATCGAGGACCTCATATCTTACGTTAAAGACAGCCGG
AATCAAGCAGTAATAATACCATGACAATCAACACCAGTACGGGGATGGAGAGAATAAA
TCCACTATCAGCTCCAGTCGGGTAAGTCTGAGGTTCACAAGCGTTCTCGTTTTGATAGC
AGCAGGTAAAAATGCTGGAAAATGTATTGGGTGCTACTGTTTCCCGAGAAGTATCACAT
CACATCATATAGCAGATCTACTATTGTTTGGAGTTAAGGAGAGTAGATTTAAAAGGCA
>Aswhite-sgRNA1-mutation 4 (WT females × mutant males)
ACGGCAGCGGTTGGAGACGCCCAGTGCAGAGCTGCTGGGGAAAAGTGCAGAAGCAA
ATAACCGATTTCTGCGGTTCATAACCGAAGCGCTACAAGGTTAAAAAGTGACTTGTAGC
TCCGTTTGCGAACGGGTCAGTCATCGAGGACCTCATATCTTACGTTAAAGACAGCCGG
AATCAAGCAGTAATAATACCATGACAATCAACACCGAACCAGTACGGGGATGGAGAGA
ATAAATCCACTATCAGCTCCAGTCGGGTAAGTCTGAGGTTCACAAGCGTTCTCGTTTTG
ATAGCAGCAGGTAAAAATGCTGGAAAATGTATTGGGTGCTACTGTTTCCCGAGAAGTAT
CACATCACATCATATAGCAGATCTACTATTGTTTGGAGTTAAGGAGAGTAGATTTAAAAG
GCA

>Aswhite-sgRNA1-mutation 5 (WT females × mutant males)
ACGGCAGCGGTTGGAGACGCCCAGTGCAGAGCTGCTGGGGAAAAGTGCAGAAGCAA
ATAACCGATTTCTGCGGTTCATAACCGAAGCGCTACAAGGTTAAAAAGTGACTTGTAGC
TCCGTTTGCGAACGGGTCAGTCATCGAGGACCTCATATCTTACGTTAAAGACAGCCGG
AATCAAGCAGTAATAATACCATGACAATCAACATGTACGGGATGGAGAGAATAAATCCA
CTATCAGCTCCAGTCGGGTAAGTCTGAGGTTCACAAGCGTTCTCGTTTTGATAGCAGCA
GGTAAAAATGCTGGAAAATGTATTGGGTGCTACTGTTTCCCGAGAAGTATCACATCACA
TCATATAGCAGATCTACTATTGTTTGGAGTTAAGGAGAGTAGATTTAAAAGGCA
>Aswhite-sgRNA2-WT
AGCGAGTAGATTGCACCAGCGTTAACAGCAGATGCTTGTTTGCTTCCTATACCCGCTAA
CAGCGCTACAGCACGAGCAGCTACCAGGACCAATCGTTGGAAGACGATGGCATCAAC
ACAACGCTGACGAACGATAAGGCGACGCTGATACAGGTGTGGAAGCCGAAGAGCTAC
GGCTCGGTGAAAGGGCAGATACCGCAGCACGAGCGGCTCACGTACACCTGGAAGGAG
ATCGACGTGTTCGGCGAGGCGCCGACCGACACGAAATCGCGCGAGCCGTTCTGCAGC
CGGCTGCGCCACTGCTTTACCTCGCGCCAGCGCCGGGACTTTAACCCGCGGAAGCACC
TGCTGAAGAATGTGACCGGTGTCGCCCGGAGCGGCGAGCTGCTGGCCGTGATGGGCA
GCTCCGGTGCGGGCAAGACGA
>Aswhite-sgRNA2-mutation 1 (WT females × WT males)
AGCGAGTAGATTGCACCAGCGTTAACAGCAGATGCTTGTTTGCTTCCTATACCCGCTAA
CAGCGCTACAGCACGAGCAGCTACCAGGACCAATCGTTGGAAGACGATGGCATCAAC
ACAACGCTGACGAACGATAAGGCGACGCTGATACAGGTGTGGAAGCCGAAGAGCTAC
GGCTCGGTGAAAGGGCAGATACCGCAGCACGAGCGGCTCACGGAAGGAGATCGACGT
GTTCGGCGAGGCGCCGACCGACACGAAATCGCGCGAGCCGTTCTGCAGCCGGCTGCG
CCACTGCTTTACCTCGCGCCAGCGCCGGGACTTTAACCCGCGGAAGCACCTGCTGAAG
AATGTGACCGGTGTCGCCCGGAGCGGCGAGCTGCTGGCCGTGATGGGCAGCTCCGGT
GCGGGCAAGACGA
>Aswhite-sgRNA2-mutation 2 (WT females × WT males)
AGCGAGTAGATTGCACCAGCGTTAACAGCAGATGCTTGTTTGCTTCCTATACCCGCTAA
CAGCGCTACAGCACGAGCAGCTACCAGGACCAATCGTTGGAAGACGATGGCATCAAC
ACAACGCTGACGAACGATAAGGCGACGCTGATACAGGTGTGGAAGCCGAAGAGCTAC
GGCTCGGTGAAAGGGCAGATACCGCAGCACGAGCGGCTCACGTAAGGAAGGAGATCG
ACGTGTTCGGCGAGGCGCCGACCGACACGAAATCGCGCGAGCCGTTCTGCAGCCGGC
TGCGCCACTGCTTTACCTCGCGCCAGCGCCGGGACTTTAACCCGCGGAAGCACCTGCT
GAAGAATGTGACCGGTGTCGCCCGGAGCGGCGAGCTGCTGGCCGTGATGGGCAGCTC
CGGTGCGGGCAAGACGA
>Aswhite-sgRNA2-mutation 3 (WT females × WT males)
AGCGAGTAGATTGCACCAGCGTTAACAGCAGATGCTTGTTTGCTTCCTATACCCGCTAA
CAGCGCTACAGCACGAGCAGCTACCAGGACCAATCGTTGGAAGACGATGGCATCAAC
ACAACGCTGACGAACGATAAGGCGACGCTGATACAGGTGTGGAAGCCGAAGAGCTAC
GGCTCGGTGAAAGGGCAGATACCGCAGCACGAGCGGCTCACGTACACCGTACGGAAG
GAGATCGACGTGTTCGGCGAGGCGCCGACCGACACGAAATCGCGCGAGCCGTTCTGC
AGCCGGCTGCGCCACTGCTTTACCTCGCGCCAGCGCCGGGACTTTAACCCGCGGAAGC
ACCTGCTGAAGAATGTGACCGGTGTCGCCCGGAGCGGCGAGCTGCTGGCCGTGATGG
GCAGCTCCGGTGCGGGCAAGACGA

>Aswhite-sgRNA2-mutation 4 (WT females × WT males)
AGCGAGTAGATTGCACCAGCGTTAACAGCAGATGCTTGTTTGCTTCCTATACCCGCTAA
CAGCGCTACAGCACGAGCAGCTACCAGGACCAATCGTTGGAAGACGATGGCATCAAC
ACAACGCTGACGAACGATAAGGCGACGCTGATACAGGTGTGGAAGCCGAAGAGCTAC
GGCTCGGTGAAAGGGCAGATACCGCAGCACGAGCGGCTCACGTACAAGGAGATCGAC
GTGTTCGGCGAGGCGCCGACCGACACGAAATCGCGCGAGCCGTTCTGCAGCCGGCTG
CGCCACTGCTTTACCTCGCGCCAGCGCCGGGACTTTAACCCGCGGAAGCACCTGCTGA
AGAATGTGACCGGTGTCGCCCGGAGCGGCGAGCTGCTGGCCGTGATGGGCAGCTCCG
GTGCGGGCAAGACGA
>Aswhite-sgRNA2-mutation 5 (WT females × WT males)
AGCGAGTAGATTGCACCAGCGTTAACAGCAGATGCTTGTTTGCTTCCTATACCCGCTAA
CAGCGCTACAGCACGAGCAGCTACCAGGACCAATCGTTGGAAGACGATGGCATCAAC
ACAACGCTGACGAACGATAAGGCGACGCTGATACAGGTGTGGAAGCCGAAGAGCTAC
GGCTCGGTGAAAGGGCAGATACCGCAGCACGAGCGGCTCACGTACACGGAAGGAGAT
CGACGTGTTCGGCGAGGCGCCGACCGACACGAAATCGCGCGAGCCGTTCTGCAGCCG
GCTGCGCCACTGCTTTACCTCGCGCCAGCGCCGGGACTTTAACCCGCGGAAGCACCTG
CTGAAGAATGTGACCGGTGTCGCCCGGAGCGGCGAGCTGCTGGCCGTGATGGGCAGC
TCCGGTGCGGGCAAGACGA
>Aswhite-sgRNA2-mutation 1 (WT females × mutant males)
AGCGAGTAGATTGCACCAGCGTTAACAGCAGATGCTTGTTTGCTTCCTATACCCGCTAA
CAGCGCTACAGCACGAGCAGCTACCAGGACCAATCGTTGGAAGACGATGGCATCAAC
ACAACGCTGACGAACGATAAGGCGACGCTGATACAGGTGTGGAAGCCGAAGAGCTAC
GGCTCGGTGAAAGGGCAGATACCGCAGCACGAGCGGCTCACGTACATCAAGGAGATC
GACGTGTTCGGCGAGGCGCCGACCGACACGAAATCGCGCGAGCCGTTCTGCAGCCGG
CTGCGCCACTGCTTTACCTCGCGCCAGCGCCGGGACTTTAACCCGCGGAAGCACCTGC
TGAAGAATGTGACCGGTGTCGCCCGGAGCGGCGAGCTGCTGGCCGTGATGGGCAGCT
CCGGTGCGGGCAAGACGA
>Aswhite-sgRNA2-mutation 2 (WT females × mutant males)
AGCGAGTAGATTGCACCAGCGTTAACAGCAGATGCTTGTTTGCTTCCTATACCCGCTAA
CAGCGCTACAGCACGAGCAGCTACCAGGACCAATCGTTGGAAGACGATGGCATCAAC
ACAACGCTGACGAACGATAAGGCGACGCTGATACAGGTGTGGAAGCCGAAGAGCTAC
GGCTCGGTGAAAGGGCAGATACCGCAGCACGAGCGGCTCACGTACACGAAGGAGATC
GACGTGTTCGGCGAGGCGCCGACCGACACGAAATCGCGCGAGCCGTTCTGCAGCCGG
CTGCGCCACTGCTTTACCTCGCGCCAGCGCCGGGACTTTAACCCGCGGAAGCACCTGC
TGAAGAATGTGACCGGTGTCGCCCGGAGCGGCGAGCTGCTGGCCGTGATGGGCAGCT
CCGGTGCGGGCAAGACGA
>Aswhite-sgRNA2-mutation 3 (WT females × mutant males)
AGCGAGTAGATTGCACCAGCGTTAACAGCAGATGCTTGTTTGCTTCCTATACCCGCTAA
CAGCGCTACAGCACGAGCAGCTACCAGGACCAATCGTTGGAAGACGATGGCATCAAC
ACAACGCTGACGAACGATAAGGCGACGCTGATACAGGTGTGGAAGCCGAAGAGCTAC
GGCTCGGTGAAAGGGCAGATACCGCAGCACGAGCGGCTCACGTACACCTTCGAAGGA
GATCGACGTGTTCGGCGAGGCGCCGACCGACACGAAATCGCGCGAGCCGTTCTGCAG
CCGGCTGCGCCACTGCTTTACCTCGCGCCAGCGCCGGGACTTTAACCCGCGGAAGCAC
CTGCTGAAGAATGTGACCGGTGTCGCCCGGAGCGGCGAGCTGCTGGCCGTGATGGGC

AGCTCCGGTGCGGGCAAGACGA

>Aswhite-sgRNA2-mutation 4 (WT females × mutant males)

AGCGAGTAGATTGCACCAGCGTTAACAGCAGATGCTTGTTTGCTTCCTATACCCGCTAA
CAGCGCTACAGCACGAGCAGCTACCAGGACCAATCGTTGGAAGACGATGGCATCAAC
ACAACGCTGACGAACGATAAGGCGACGCTGATACAGGTGTGGAAGCCGAAGAGCTAC
GGCTCGGTGAAAGGGCAGATACCGCAGCACGAGCGGCTCACGGCGGAAGGAGATCGA
CGTGTTCGGCGAGGCGCCGACCGACACGAAATCGCGCGAGCCGTTCTGCAGCCGGCT
GCGCCACTGCTTTACCTCGCGCCAGCGCCGGGACTTTAACCCGCGGAAGCACCTGCTG
AAGAATGTGACCGGTGTCGCCCGGAGCGGCGAGCTGCTGGCCGTGATGGGCAGCTCC
GGTGCGGGCAAGACGA

>Aswhite-sgRNA2-mutation 5 (WT females × mutant males)

AGCGAGTAGATTGCACCAGCGTTAACAGCAGATGCTTGTTTGCTTCCTATACCCGCTAA
CAGCGCTACAGCACGAGCAGCTACCAGGACCAATCGTTGGAAGACGATGGCATCAAC
ACAACGCTGACGAACGATAAGGCGACGCTGATACAGGTGTGGAAGCCGAAGAGCTAC
GGCTCGGTGAAAGGGCAGATACCGCAGCACGAGCGGCTCACGTACACCACGGGAAGG
AGATCGACGTGTTCGGCGAGGCGCCGACCGACACGAAATCGCGCGAGCCGTTCTGCA
GCCGGCTGCGCCACTGCTTTACCTCGCGCCAGCGCCGGGACTTTAACCCGCGGAAGCA
CCTGCTGAAGAATGTGACCGGTGTCGCCCGGAGCGGCGAGCTGCTGGCCGTGATGGG
CAGCTCCGGTGCGGGCAAGACGA

>Asyellow-sgRNA1-WT

TACAGCTCCTGGCTGGAAGTGCGTGTGTCTTGCGGAGGCGTGTGTATAGCGAAGCCAT
GCAGAGAACGATGTACGGTTTGCTGATCGCGGTGTGCCTGGCCGCCGGTAGTGTCAGT
GCGACGCAGAAGCTGCAGGAGCGCTACAGCTGGCAGCAGCTCGACTTCGTCTTCCCC
AATCAACGCCTCAAGCAACAGGCCCTGGCTAGCGGGGACTACGTCCCGACCAACGGT
CTGCCGGTCGGCATCGAGCGCTGGGAGAATAAGCTGTTCGTGTCTGTGCCGAGATGGA
AGGATGGTACGTATGCACGCCTTGGTGGCAGACTTCTCGAGTGACCGTATCGTACAGTT
CAACCGATCTCGTGCGCCTCTACTTAGTGTCCCCAAAATTCTCAATCCCCGCGTGACAG
ATATTCGCGAGTAGTGCTCCCAAAATTCCCAGATTCGTCCCGTTAT

>Asyellow-sgRNA1-mutation 1 (WT females × WT males)

TACAGCTCCTGGCTGGAAGTGCGTGTGTCTTGCGGAGGCGTGTGTATAGCGAAGCCAT
GCAGAGAACGATGTACGGTTTGCTGATCGCGGTGTGCCTGGCCGCCGGTAGTGTCAGT
GCGACGCAGAAGCTGCAGGAGCGCTACAGCTGGCAGCAGCTCGACTTCGTCTTCCCC
AATCAACGCCTCAAGCAACAGGCCCTGGCTAGCGGGGACTACGTCCCGACCAACCGG
CTACGACCGAAGGCCGGTCGGCATCGAGCGCTGGGAGAATAAGCTGTTCGTGTCTGTG
CCGAGATGGAAGGATGGTACGTATGCACGCCTTGGTGGCAGACTTCTCGAGTGACCGT
ATCGTACAGTTCAACCGATCTCGTGCGCCTCTACTTAGTGTCCCCAAAATTCTCAATCC
CCGCGTGACAGATATTCGCGAGTAGTGCTCCCAAAATTCCCAGATTCGTCCCGTTAT

>Asyellow-sgRNA1-mutation 2 (WT females × WT males)

TACAGCTCCTGGCTGGAAGTGCGTGTGTCTTGCGGAGGCGTGTGTATAGCGAAGCCAT
GCAGAGAACGATGTACGGTTTGCTGATCGCGGTGTGCCTGGCCGCCGGTAGTGTCAGT
GCGACGCAGAAGCTGCAGGAGCGCTACAGCTGGCAGCAGCTCGACTTCGTCTTCCCC
AATCAACGCCTCAAGCAACAGGCCCTGGCTAGCGGGGACTACGTCCCGACCAACGGT
CGATGCCGGTCGGCATCGAGCGCTGGGAGAATAAGCTGTTCGTGTCTGTGCCGAGATG
GAAGGATGGTACGTATGCACGCCTTGGTGGCAGACTTCTCGAGTGACCGTATCGTACA

GTTCAACCGATCTCGTGCGCCTCTACTTAGTGTCCCCAAAATTCTCAATCCCCGCGTGA
CAGATATTCGCGAGTAGTGCTCCCAAAATTCCCAGATTCGTCCCGTTAT
>Asyellow-sgRNA1-mutation 3 (WT females × WT males)
TACAGCTCCTGGCTGGAAGTGCGTGTGTCTTGCGGAGGCGTGTGTATAGCGAAGCCAT
GCAGAGAACGATGTACGGTTTGCTGATCGCGGTGTGCCTGGCCGCCGGTAGTGTCAGT
GCGACGCAGAAGCTGCAGGAGCGCTACAGCTGGCAGCAGCTCGACTTCGTCTTCCCC
AATCAACGCCTCAAGCAACAGGCCCTGGCTAGCGGGGACTACGTCCCGACCAACGGT
CGAACGGCCGGTCGGCATCGAGCGCTGGGAGAATAAGCTGTTCGTGTCTGTGCCGAG
ATGGAAGGATGGTACGTATGCACGCCTTGGTGGCAGACTTCTCGAGTGACCGTATCGTA
CAGTTCAACCGATCTCGTGCGCCTCTACTTAGTGTCCCCAAAATTCTCAATCCCCGCGT
GACAGATATTCGCGAGTAGTGCTCCCAAAATTCCCAGATTCGTCCCGTTAT
>Asyellow-sgRNA1-mutation 4 (WT females × WT males)
TACAGCTCCTGGCTGGAAGTGCGTGTGTCTTGCGGAGGCGTGTGTATAGCGAAGCCAT
GCAGAGAACGATGTACGGTTTGCTGATCGCGGTGTGCCTGGCCGCCGGTAGTGTCAGT
GCGACGCAGAAGCTGCAGGAGCGCTACAGCTGGCAGCAGCTCGACTTCGTCTTCCCC
AATCAACGCCTCAAGCAACAGGCCCTGGCTAGCGGGGACTACGTCCCGACCAACGGT
CGGCATCGAGCGCTGGGAGAATAAGCTGTTCGTGTCTGTGCCGAGATGGAAGGATGGT
ACGTATGCACGCCTTGGTGGCAGACTTCTCGAGTGACCGTATCGTACAGTTCAACCGAT
CTCGTGCGCCTCTACTTAGTGTCCCCAAAATTCTCAATCCCCGCGTGACAGATATTCGC
GAGTAGTGCTCCCAAAATTCCCAGATTCGTCCCGTTAT
>Asyellow-sgRNA1-mutation 1 (WT females × mutant males)
TACAGCTCCTGGCTGGAAGTGCGTGTGTCTTGCGGAGGCGTGTGTATAGCGAAGCCAT
GCAGAGAACGATGTACGGTTTGCTGATCGCGGTGTGCCTGGCCGCCGGTAGTGTCAGT
GCGACGCAGAAGCTGCAGGAGCGCTACAGCTGGCAGCAGCTCGACTTCGTCTTCCCC
AATCAACGCCTCAAGCAACAGGCCCTGGCTAGCGGGGACTACGTCCCGACCAACGCT
CGGCATCGAGCGCTGGGAGAATAAGCTGTTCGTGTCTGTGCCGAGATGGAAGGATGGT
ACGTATGCACGCCTTGGTGGCAGACTTCTCGAGTGACCGTATCGTACAGTTCAACCGAT
CTCGTGCGCCTCTACTTAGTGTCCCCAAAATTCTCAATCCCCGCGTGACAGATATTCGC
GAGTAGTGCTCCCAAAATTCCCAGATTCGTCCCGTTAT
>Asyellow-sgRNA1-mutation 2 (WT females × mutant males)
TACAGCTCCTGGCTGGAAGTGCGTGTGTCTTGCGGAGGCGTGTGTATAGCGAAGCCAT
GCAGAGAACGATGTACGGTTTGCTGATCGCGGTGTGCCTGGCCGCCGGTAGTGTCAGT
GCGACGCAGAAGCTGCAGGAGCGCTACAGCTGGCAGCAGCTCGACTTCGTCTTCCCC
AATCAACGCCTCAAGCAACAGGCCCTGGCTAGCGGGGACTACGTTCCGACCAACGGT
CGAACGCTGGGAGAATAAGCTGTTCGTGTCTGTGCCGAGATGGAAGGATGGTACGTAT
GCACGCCTTGGTGGCAGACTTCTCGAGTGACCGTATCGTACAGTTCAACCGATCTCGT
GCGCCTCTACTTAGTGTCCCCAAAATTCTCAATCCCCGCGTGACAGATATTCGCGAGTA
GTGCTCCCAAAATTCCCAGATTCGTCCCGTTAT
>Asyellow-sgRNA1-mutation 3 (WT females × mutant males)
TACAGCTCCTGGCTGGAAGTGCGTGTGTCTTGCGGAGGCGTGTGTATAGCGAAGCCAT
GCAGAGAACGATGTACGGTTTGCTGATCGCGGTGTGCCTGGCCGCCGGTAGTGTCAGT
GCGACGCAGAAGCTGCAGGAGCGCTACAGCTGGCAGCAGCTCGACTTCGTCTTCCCC
AATCAACGCCTCAAGCAACAGGCCCTGGCTAGCGGGGACTACGTCCCGACCAACCAC
CGGTCGGCATCGAGCGCTGGGAGAATAAGCTGTTCGTGTCTGTGCCGAGATGGAAGGA

TGGTACGTATGCACGCCTTGGTGGCAGACTTCTCGAGTGACCGTATCGTACAGTTCAAC
CGATCTCGTGCGCCTCTACTTAGTGTCCCCAAAATTCTCAATCCCCGCGTGACAGATATT
CGCGAGTAGTGCTCCCAAAATTCCCAGATTCGTCCCGTTAT
>Asyellow-sgRNA1-mutation 4 (WT females × mutant males)
TACAGCTCCTGGCTGGAAGTGCGTGTGTCTTGCGGAGGCGTGTGTATAGCGAAGCCAT
GCAGAGAACGATGTACGGTTTGCTGATCGCGGTGTGCCTGGCCGCCGGTAGTGTCAGT
GCGACGCAGAAGCTGCAGGAGCGCTACAGCTGGCAGCAGCTCGACTTCGTCTTCCCC
AATCAACGCCTCAAGCAACAGGCCCTGGCTAGCGGGGACTACGTCCCGACCAACGGT
CGAAATCGAGCGCTGGGAGAATAAGCTGTTCGTGTCTGTGCCGAGATGGAAGGATGGT
ACGTATGCACGCCTTGGTGGCAGACTTCTCGAGTGACCGTATCGTACAGTTCAACCGAT
CTCGTGCGCCTCTACTTAGTGTCCCCAAAATTCTCAATCCCCGCGTGACAGATATTCGC
GAGTAGTGCTCCCAAAATTCCCAGATTCGTCCCGTTAT
>YELLOW-sgRNA1-mutation 5 (WT females × mutant males)
TACAGCTCCTGGCTGGAAGTGCGTGTGTCTTGCGGAGGCGTGTGTATAGCGAAGCCAT
GCAGAGAACGATGTACGGTTTGCTGATCGCGGTGTGCCTGGCCGCCGGTAGTGTCAGT
GCGACGCAGAAGCTGCAGGAGCGCTACAGCTGGCAGCAGCTCGACTTCGTCTTCCCC
AATCAACGCCTCAAGCAACAGGCCCTGGCTAGCGGGGACTACGTCCCGACCAACGGT
CGGTCGGCATCGAGCGCTGGGAGAATAAGCTGTTCGTGTCTGTGCCGAGATGGAAGGA
TGGTACGTATGCACGCCTTGGTGACAGACTTCTCGAGTGACCGTATCGTACAGTTCAAC
CGATCTCGTGCGCCTCTACTTAGTGTCCCCAAAATTCTCAATCCCCGCGTGACAGATATT
CGCGAGTAGTGCTCCCAAAATTCCCAGATTCGTCCCGTTAT
>Asyellow-sgRNA2-WT
GACGCCTGTGGGCCCTGGACACCGGAACCGTCGGTATCGGTAACACCACCCAGCAGC
TGTGCCCGTACGCGCTCAACGTGTGGGACCTGAAGACGAACCGCCGCATCCGCCGCTA
CGAGCTGCGCCCAGAAGACACCAACCCGAACACGTTCATCGCCAACATCGCCATCGA
CATGGGTCGCAGCTGCGACGACACGTTTGCGTACATGTCCGACGAACTGGGCTACGGT
CTGATCGTGTACTCGTTCGAGCAGAACAAGTCGTGGCGGTTCGCTCACAGCTTCTTCTT
CCCGGATCCGCTGCGCGGTGACTTCAACGTTGCCGGCCTGAACTTCCAGTGGGGCGAG
GAGGGCATCTTCGGCATGTCGTTGACGCCGCTGCAAGCCGATGGCTACCGGACGCTCT
ACTTCTCGCCGCTTGCCAGCCATCGCGAGTTCATG
>Asyellow-sgRNA2-mutation 1 (WT females × WT males)
GACGCCTGTGGGCCCTGGACACCGGAACCGTCGGTATCGGTAACACCACCCAGCAGC
TGTGCCCGTACGCGCTCAACGTGTGGGACCTGAAGACGAACCGCCGCATCCGCCGCTA
CGAGCTGCGCCCAGAAGACACCAACCCGAACACGTTCATCGCCAACATCGCCATCGA
CATGGGTCGCAGCTGCGACGACACGTTTGCGTACATGTCCGACGAACTGGCAGAAATA
TCAACAAACGGTCTGATCGTGTACTCGTTCGAGCAGAACAAGTCGTGGCGGTTCGCTC
ACAGCTTCTTCTTCCCGGATCCGCTGCGCGGTGACTTCAACGTTGCCGGCCTGAACTT
CCAGTGGGGCGAGGAGGGCATCTTCGGCATGTCGTTGACGCCGCTGCAAGCCGATGG
CTACCGGACGCTCTACTTCTCGCCGCTTGCCAGCCATCGCGAGTTCATG
>Asyellow-sgRNA2-mutation 2 (WT females × WT males)
GACGCCTGTGGGCCCTGGACACCGGAACCGTCGGTATCGGTAACACCACCCAGCAGC
TGTGCCCGTACGCGCTCAACGTGTGGGACCTGAAGACGAACCGCCGCATCCGCCGCTA
CGAGCTGCGCCCAGAAGACACCAACCCGAACACGTTCATCGCCAACATCGCCATCGA
CATGGGTCGCAGCTGCGACGACACGTTTGCGTACATGTCCGACGAACGACGGTCTGAT

CGTGTACTCGTTCGAGCAGAACAAGTCGTGGCGGTTCGCTCACAGCTTCTTCTTCCCG
GATCCGCTGCGCGGTGACTTCAACGTTGCCGGCCTGAACTTCCAGTGGGGCGAGGAG
GGCATCTTCGGCATGTCGTTGACGCCGCTGCAAGCCGATGGCTACCGGACGCTCTACTT
CTCGCCGCTTGCCAGCCATCGCGAGTTCATG
>Asyellow-sgRNA2-mutation 3 (WT females × WT males)
GACGCCTGTGGGCCCTGGACACCGGAACCGTCGGTATCGGTAACACCACCCAGCAGC
TGTGCCCGTACGCGCTCAACGTGTGGGACCTGAAGACGAACCGCCGCATCCGCCGCTA
CGAGCTGCGCCCAGAAGACACCAACCCGAACACGTTCATCGCCAACATCGCCATCGA
CATGGGTCGCAGCTGCGACGACACGTTTGCGTACATGTCCGACGAACTACGGTCTGAT
CGTGTACTCGTTCGAGCAGAACAAGTCGTGGCGGTTCGCTCACAGCTTCTTCTTCCCG
GATCCGCTGCGCGGTGACTTCAACGTTGCCGGCCTGAACTTCCAGTGGGGCGAGGAG
GGCATCTTCGGCATGTCGTTGACGCCGCTGCAAGCCGATGGCTACCGGACGCTCTACTT
CTCGCCGCTTGCCAGCCATCGCGAGTTCATG
>Asyellow-sgRNA2-mutation 4 (WT females × WT males)
GACGCCTGTGGGCCCTGGACACCGGAACCGTCGGTATCGGTAACACCACCCAGCAGC
TGTGCCCGTACGCGCTCAACGTGTGGGACCTGAAGACGAACCGCCGCATCCGCCGCTA
CGAGCTGCGCCCAGAAGACACCAACCCGAACACGTTCATCGCCAACATCGCCATCGA
CATGGGTCGCAGCTGCGACGACACGTTTGCGTACATGTCCGACGAACTGGTCTGATCG
TGTACTCGTTCGAGCAGAACAAGTCGTGGCGGTTCGCTCACAGCTTCTTCTTCCCGGA
TCCGCTGCGCGGTGACTTCAACGTTGCCGGCCTGAACTTCCAGTGGGGCGAGGAGGG
CATCTTCGGCATGTCGTTGACGCCGCTGCAAGCCGATGGCTACCGGACGCTCTACTTCT
CGCCGCTTGCCAGCCATCGCGAGTTCATG
>Asyellow-sgRNA2-mutation 5 (WT females × WT males)
GACGCCTGTGGGCCCTGGACACCGGAACCGTCGGTATCGGTAACACCACCCAGCAGC
TGTGCCCGTACGCGCTCAACGTGTGGGACCTGAAGACGAACCGCCGCATCCGCCGCTA
CGAGCTGCGCCCAGAAGACACCAACCCGAACACGTTCATCGCCAACATCGCCATCGA
CATGGGTCGCAGCTGCGACGACACGTTTGCGTACATGTCCGACGAACTGGGTCTACGG
TCTGATCGTGTACTCGTTCGAGCAGAACAAGTCGTGGCGGTTCGCTCACAGCTTCTTCT
TCCCGGATCCGCTGCGCGGTGACTTCAACGTTGCCGGCCTGAACTTCCAGTGGGGCGA
GGAGGGCATCTTCGGCATGTCGTTGACGCCGCTGCAAGCCGATGGCTACCGGACGCTC
TACTTCTCGCCGCTTGCCAGCCATCGCGAGTTCATG
>Asyellow-sgRNA2-mutation 1 (WT females × mutant males)
GACGCCTGTGGGCCCTGGACACCGGAACCGTCGGTATCGGTAACACCACCCAGCAGC
TGTGCCCGTACGCGCTCAACGTGTGGGACCTGAAGACGAACCGCCGCATCCGCCGCTA
CGAGCTGCGCCCAGAAGACACCAACCCGAACACGTTCATCGCCAACATCGCCATCGA
CATGGGTCGCAGCTGCGACGACACGTTTGCGTACATGTCCGACGAACTGGTGGGCACG
ACGGTCTGATCGTGTACTCGTTCGAGCAGAACAAGTCGTGGCGGTTCGCTCACAGCTT
CTTCTTCCCGGATCCGCTGCGCGGTGACTTCAACGTTGCCGGCCTGAACTTCCAGTGG
GGCGAGGAGGGCATCTTCGGCATGTCGTTGACGCCGCTGCAAGCCGATGGCTACCGGA
CGCTCTACTTCTCGCCGCTTGCCAGCCATCGCGAGTTCATG
>Asyellow-sgRNA2-mutation 2 (WT females × mutant males)
GACGCCTGTGGGCCCTGGACACCGGAACCGTCGGTATCGGTAACACCACCCAGCAGC
TGTGCCCGTACGCGCTCAACGTGTGGGACCTGAAGACGAACCGCCGCATCCGCCGCTA
CGAGCTGCGCCCAGAAGACACCAACCCGAACACGTTCATCGCCAACATCGCCATCGA

CATGGGTCGCAGCTGCGACGACACGTTTGCGTACATGTCCGACGAACTGGGTCGTGTA
CTCGTTCGAGCAGAACAAGTCGTGGCGGTTCGCTCACAGCTTCTTCTTCCCGGATCCG
CTGCGCGGTGACTTCAACGTTGCCGGCCTGAACTTCCAGTGGGGCGAGGAGGGCATC
TTCGGCATGTCGTTGACGCCGCTGCAAGCCGATGGCTACCGGACGCTCTACTTCTCGCC
GCTTGCCAGCCATCGCGAGTTCATG

>Asyellow-sgRNA2-mutation 3 (WT females × mutant males)

GACGCCTGTGGGCCCTGGACACCGGAACCGTCGGTATCGGTAACACCACCCAGCAGC
TGTGCCCGTACGCGCTCAACGTGTGGGACCTGAAGACGAACCGCCGCATCCGCCGCTA
CGAGCTGCGCCCAGAAGACACCAACCCGAACACGTTCATCGCCAACATCGCCATCGA
CATGGGTCGCAGCTGCGACGACACGTTTGCGTACATGTCCGACGAACTGGGTCTGATC
GTGTACTCGTTCGAGCAGAACAAGTCGTGGCGGTTCGCTCACAGCTTCTTCTTCCCGG
ATCCGCTGCGCGGTGACTTCAACGTTGCCGGCCTGAACTTCCAGTGGGGCGAGGAGG
GCATCTTCGGCATGTCGTTGACGCCGCTGCAAGCCGATGGCTACCGGACGCTCTACTTC
TCGCCGCTTGCCAGCCATCGCGAGTTCATG

>Asyellow-sgRNA2-mutation 4 (WT females × mutant males)

GACGCCTGTGGGCCCTGGACACCGGAACCGTCGGTATCGGTAACACCACCCAGCAGC
TGTGCCCGTACGCGCTCAACGTGTGGGACCTGAAGACGAACCGCCGCATCCGCCGCTA
CGAGCTGCGCCCAGAAGACACCAACCCGAACACGTTCATCGCCAACATCGCCATCGA
CATGGGTCGCAGCTGCGACGACACGTTTGCGTACATGTCCGACGAATGTACTCGTTCG
AGCAGAACAAGTCGTGGCGGTTCGCTCACAGCTTCTTCTTCCCGGATCCGCTGCGCGG
TGACTTCAACGTTGCCGGCCTGAACTTCCAGTGGGGCGAGGAGGGCATCTTCGGCATG
TCGTTGACGCCGCTGCAAGCCGATGGCTACCGGACGCTCTACTTCTCGCCGCTTGCCA
GCCATCGCGAGTTCATG

>Asyellow-sgRNA2-mutation 5 (WT females × mutant males)

GACGCCTGTGGGCCCTGGACACCGGAACCGTCGGTATCGGTAACACCACCCAGCAGC
TGTGCCCGTACGCGCTCAACGTGTGGGACCTGAAGACGAACCGCCGCATCCGCCGCTA
CGAGCTGCGCCCAGAAGACACCAACCCGAACACGTTCATCGCCAACATCGCCATCGA
CATGGGTCGCAGCTGCGACGACACGTTTGCGTACATGTCCGACGGTACGTACCGGTCT
GATCGTGTACTCGTTCGAGCAGAACAAGTCGTGGCGGTTCGCTCACAGCTTCTTCTTC
CCGGATCCGCTGCGCGGTGACTTCAACGTTGCCGGCCTGAACTTCCAGTGGGGCGAG
GAGGGCATCTTCGGCATGTCGTTGACGCCGCTGCAAGCCGATGGCTACCGGACGCTCT
ACTTCTCGCCGCTTGCCAGCCATCGCGAGTTCATG
