## Supplementary material for "High-efficiency gene editing in *Anopheles sinensis* using ReMOT control": 20230828 Supplemental materials and methods.pdf

1. Mosquitoes

The WT strain used for adult injection is the LS-WX homozygous strain (inbreeding for over 40 generations). The rearing conditions were as follows: 27-29°C, 75% ± 5% relative humidity, and a 12:12 h (light: dark) photoperiod. Larvae are reared in plastic box containing dechlorinated tap water and fed micro pellets fish food. After pupation, they are transferred to white plastic bucket covered with gauze until eclosion. The adults are fed with a 10% glucose solution. Approximately one day after eclosion, females and males are allowed to mate extensively (approximately 72 hours). Subsequently, female mosquitoes are subjected to a 10 h period of starvation before being provided a blood feeding.

2. Expression pattern of *AsVg* transcripts

Female mosquitoes at different stages (0 h, 6 h, 12 h, 18 h, 24 h, 30 h) post blood feeding (PBF) were selected. Total RNA was extracted using the Trizol method and cDNA synthesis of the samples was performed using the PrimeScript™ RT reagent Kit with gDNA Eraser (Perfect Real Time) kit (Takara, Japan). Real time quantitative PCR (RT-qPCR) primers were designed (Table S1) based on the reference sequence of the *AsVg* gene (gene ID: *Sinensis_GLEAN_10014898*, obtained from unpublished data of the *Anopheles sinensis* genome). *AsRPL49* was used as the internal reference gene. The protocol for qPCR was conducted following previously published methods (Qiao et al., 2016). The $2^{-\Delta Ct}$ method was employed to calculate the relative expression level of the *AsVg*.

3. Plasmids Construction

The expression vector *pET28a* was stored in our laboratory. The coding sequence of the P2C delivering peptide and the Linker fragment sequences were synthesized separately by Beijing Qingke Biotechnology Co., Ltd. and cloned into the *pMD19-T Simple* vector. The fragment of the enhanced red fluorescent protein gene *DsRed* was amplified from the *pBac-3×P3-DsRed* plasmid stored in our laboratory, and the *Cas9* fragment was amplified from the *pBac-NOS-Cas9* plasmid provided by Dr. Yun Wang from Chongqing University.

The above fragments were amplified using PrimeSTAR® Max DNA Polymerase (Takara, Japan). The *pET28a* plasmid was digested with QuickCut™ *Nde* I and QuickCut™ *Xho* I enzymes (Takara, Japan), respectively, and then ligated with the corresponding fragments mentioned above. Finally, the *pET28a-P2C-DsRed*, *pET28a-DsRed*, *pET28a-P2C-Linker-Cas9* and *pET28a-Cas9* recombinant protein vectors required for this study were constructed. The primer sequences required for all recombinant protein vectors are provided in Table S1.

4. Expression and purification of recombinant proteins

The successfully constructed *pET28a-P2C-DsRed*, *pET28a-DsRed*, *pET28a-P2C-Linker-Cas9* and *pET28a-Cas9* plasmids were separately transformed into *Escherichia coli* BL21(DE3) competent cells (TransGen Biotech, China). The recombinant bacteria were cultured in TB medium at 37°C until the OD600 value reached 0.6-0.8. Then, induction was performed using 0.5 mmol/L IPTG at 16°C with shaking at 180 rpm for 16-20 h. The cells were collected by centrifugation at 7,000 rpm for 15 min. The cell pellets were resuspended in Lysis Buffer (50 mM $NaH_2PO_4$, 300 mM NaCl, 10 mM imidazole, pH adjusted to 8.0) at a ratio of 1:10 (w/v), along with the addition of 1 mM PMSF. The mixture was subjected to ultrasonication for cell disruption. After centrifugation at 10,000 rpm for 30 min at 4°C, the supernatant was collected. The supernatant was filtered through a 0.45 μm membrane and purified using Ni-NTA nickel columns (Qiagen, Germany) to obtain the desired proteins, pET28a-P2C-DsRed, pET28a-DsRed, and pET28a-P2C-Linker-Cas9. Protein specificity was confirmed through polyacrylamide gel electrophoresis (SDS-PAGE) and Western blot analysis using the SDS-PAGE Gel Quick Preparation Kit (Beyotime, China). Desalting was performed by dialysis using Spectra/Por® 6 Dialysis Membranes (Sangon Biotech, China), and the protein concentration was determined using the BCA Protein Assay Kit (Beyotime, China).

In this study, the Western blot procedure was as follows: Protein samples separated by a 12% SDS-PAGE gel were transferred onto PVDF membranes (Roche, Switzerland). The membranes were blocked at room temperature for 1 h with a solution containing 5% protein powder (Sangon Biotech, China) and 10% sheep serum (Sigma, USA). After washing three times with TBST solution (Thermo, USA), the membranes were incubated overnight at 4°C with a mouse monoclonal Anti-His antibody (Roche, Switzerland) or Anti-Flag antibody (Roche, Switzerland) diluted at a ratio of 1:6,000. Following three washes with TBST solution, the membranes were incubated at room temperature for 1 h with an HRP-conjugated goat anti-mouse IgG antibody (Sigma, USA) diluted at a ratio of 1:10,000. After washing, the membranes were incubated with the ECL chemiluminescent substrate from the ECL Western Blotting Detection Kit (Thermo, USA) for 5 min. The protein bands were visualized and captured using the Universal Hood III, XR, XRS fluorescence/chemiluminescence imaging system (BIO-RAD, USA) and saved for analysis.

5. The synthesis and *in vitro* cleavage of sgRNA

Design and select the sgRNA sites of *Aswhite* (*Sinensis_GLEAN_10014141*) and *Asyellow* (*Sinensis_GLEAN_10005782*) using the sgRNA prediction website (http://crispor.tefor.net/). Two target sites were chosen for per gene editing analysis. The sgRNA was synthesized using the T7 RiboMAX™ Express RNAi System Kit (Promega, USA). The DNA sequences containing gRNA sites were amplified from the genome of *A. sinensis* using PrimeSTAR® Max DNA Polymerase (Takara, Japan) as templates for *in vitro* cleavage. The steps for *in vitro* cleavage can be summarized as follows: RNase-free water, sgRNA (approximately 120 ng/μL), and pET28a-P2C-Linker-Cas9 or pET28a -Cas9 protein (approximately 130-160 ng/μL) were gently mixed and incubated at 25°C for 20 min. Then, DNA template (approximately 160-200 ng) and 10×NEBuffer™ 3.1 (New England Biolabs, USA) were added, mixed well, and incubated at 37°C for 1 h. Finally, Proteinase K (20 mg/ml) (Jiangsu Cowin Biotech, China) was added, and the mixture was digested at 56°C for 20 minutes. The reaction products were detected by agarose gel electrophoresis. The primers required for sgRNA transcription and *in vitro* cleavage are listed in Table S1.

6. Anatomical observation of ovarian tissues

Female mosquitoes were anesthetized on ice at 20 and 40 h post blood feeding. The abdominal end was clamped with surgical forceps, and the ovarian tissues were quickly extracted and washed in cold PBS buffer (HyClone, USA). The development of the ovaries at different time post blood feeding was observed using a Research Biological Microscope BX53 fluorescence microscope (Olympus, Japan).

7. Microinjection

A borosilicate glass needle with a diameter of 0.5 mm (WPI, USA) was pulled to create a sharp and fine glass tip using the SUTTER P-2000 micropipette puller (SUTTER, USA). During injection, the mixture was loaded into the glass needle and the injection parameters of the WPI PV830 Picopump (WPI, USA) were adjusted accordingly. Female mosquitoes at 20 h PBF were anesthetized on ice, and approximately 250 nL of the injection solution was injected into the thorax lumen. After injection, the female mosquitoes were maintained following Part 1 and allowed to lay eggs.

8. Ovarian delivery and detection of pET28a-P2C-DsRed and pET28a-DsRed protein

pET28a-P2C-DsRed recombinant protein (560 ng/μL) was injected into the thorax lumen of female mosquitoes after 20 h of blood feeding. Ovaries were dissected at 40 h of post blood feeding, and the enrichment of the recombinant protein within the entire ovarian tissue was observed using Olympus BX53 fluorescence microscope. The ovarian tissues displaying red fluorescence were embedded in Tissue-Tek® O.C.T. Compound (SAKURA, USA) medium in capsule-shaped molds. After the tissue blocks were solidified at -20°C, the samples were affixed to a slice tray and sectioned using a Thermo HM525 NX Freezing Microtome (Thermo, USA) with a thickness of 4 μm. The enrichment of the recombinant protein in the sliced ovarian tissues was further observed using BX63 fluorescence microscope (Olympus, Japan). Injection of pET28a-DsRed (450 ng/μL, approximately equal molar ratio to the treatment group) recombinant protein was used as the control group.

After homogenizing the ovaries, the cells were lysed using Cell and IP Lysis Buffer (Thermo, USA) to extract proteins. The enrichment of the injected recombinant proteins was then detected using Western blot analysis.

9. Determination concentration of saponin

The recombinant protein pET28a-P2C-Linker-Cas9 was mixed with the endosomal escape reagent (EER) saponin (SIGMA, USA) to create a gradient of six injection mixtures with saponin concentrations of 100 ng/μL, 150 ng/μL, 200 ng/μL, 250 ng/μL, 300 ng/μL, and 350 ng/μL. The working concentration of the recombinant protein in each gradient was maintained at 2600 ng/μL. Following Part 7, the injection mixture was injected into the thorax lumen of female mosquitoes. pET28a-DsRed recombinant protein (2500 ng/μL, approximately equal molar ratio to the treatment group) was injected as a control group. The number of laid eggs and the hatching rate of larvae were recorded for each injected individuals in order to determine the optimum EER concentration.

10. Mutation screening and genotyping

For the two target sites of *Aswhite* and the two target sites of *Asyellow*, the treatment group was set as pET28a-P2C-Linker-Cas9 + sgRNA + Saponin, while the control group was set as pET28a-Cas9 + sgRNA + Saponin. The specific *in vitro* binding and incubation system included a working concentration of approximately 2600 ng/μL for pET28a-P2C-Linker-Cas9/pET28a-Cas9, a working concentration of approximately 1100 ng/μL for sgRNA, and a working concentration of saponin with 100 ng/μL. After incubating the Cas9 ribonucleoprotein (RNP) mixture at 25°C for 20 min, Saponin was gently added and the injection mixture was kept on ice for further use. Following Part 7, injections were administered to WT females that mate with WT males at 20 hours post-blood-feeding. For the *Aswhite* knockout treatment group, the coloring status of the compound eyes was observed and the $G_0$ gene-editing efficiency (GEF) was observed for eye coloration from the pupal stage onwards. For the *Asyellow* knockout treatment group, was observed for coloring pattern of the epidermis and sclerotized regions was observed, and GEF was recorded three days after eclosion.

Genomic DNA was extracted from individuals with mutant phenotypes using the Universal Genomic DNA Kit (Jiangsu Cowin Biotech, China). Primers were designed in the flanking regions of the *Aswhite* and *Asyellow* gRNA sites on the genome, and PCR amplification was performed. The PCR products were purified using the E.Z.N.A® Gel Extraction Kit (Omega, USA) and subsequently cloned using the pEASY®-Blunt Zero Cloning Kit (TransGen Biotech, China). Types of gene editing were analyzed through sequencing. The primers required for genotyping are listed in Table S1.

11. Sex-linked inheritance analysis

To analyze the linkage relationship between the *Aswhite* (or *Asyellow*) and X chromosome, outcross combinations of mutant females mated with WT males and mutant males mated with WT females were used, respectively. Each outcross combination consisted of 10 male and 10 female parents. The gender and phenotype of the $F_1$ generation individuals were observed and recorded. Well-developed adult individuals from the $F_1$ generation were selected for full-sib mating in a ratio of female to male of 15:15. The gender ratio and phenotype of the $F_2$ generation individuals were observed and recorded.

12. Correction of editing efficiency

For the correction of editing efficiency for *Aswhite* and *Asyellow*, injection should be performed on WT females that mated with mutant males at post blood feeding. The methods of injection, types of injection mixtures, observation and statistical analysis of offspring phenotypes after injection, and molecular detection of mutant individuals followed Part 7 and Part 10.
