## Supplementary material for "High-efficiency gene editing in *Anopheles sinensis* using ReMOT control": 20230828 Supplemental tables.pdf

Table S1. Primers used in this study

| Primer names | Primer sequences (5’-3’) | Purpose |
|---|---|---|
| *Aswhite* target site-1-F | GAAATTAATACGACTCACTATAG*TCCATCCCCGTACTGGTCAT*GTTTTAGAGCTAGAAATAGCAAG | Synthesis of *Aswhite* sgRNA |
| *Aswhite* target site-2-F | GAAATTAATACGACTCACTATAG*GCGGCTCACGTACACCTGGA*GTTTTAGAGCTAGAAATAGCAAG | |
| *Asyellow* target site-1-F | GAAATTAATACGACTCACTATAG*CGTCCCGACCAACGGTCTGC*GTTTTAGAGCTAGAAATAGCAAG | Synthesis of *Asyellow* sgRNA |
| *Asyellow* target site-2-F | GAAATTAATACGACTCACTATAG*ATGTCCGACGAACTGGGCTA*GTTTTAGAGCTAGAAATAGCAAG | |
| sgRNA-r | AAAAGCACCGACTCGGTGCCACTTTTTCAAGTTGATAACGGACTAGCCTTATTTTAACTTGCTATTTCTAGCTCTAAAAC | sgRNA downstream primer |
| *Aswhite* PCR target1 F | GGCAAAGGAATAGCGAAGAGTC | *In vitro* cleavage experiment of the *Aswhite* |
| *Aswhite* PCR target1 R | TGTTGTTGGGCTTCGTTATGC | |
| *Aswhite* PCR target2 F | AGGTGTGGAAGCCGAAGAGC | |
| *Aswhite* PCR target2 R | CGTCTGGTGGATGGTGAGGAT | |
| *Asyellow* target1-F | CGACGGTAGTGTCAGTGCGA | *In vitro* cleavage experiment of the *Asyellow* |
| *Asyellow* target1-R | CGAGAAGTCTGCCACCAAGG | |
| *Asyellow* target2-F | ATAAGTGCGGACGCCTGTG | |
| *Asyellow* target2-R | ATGCCGAAGATGCCCTCC | |
| *Aswhite* detection-w1-F | TCGGCTGAAGAAGTGAAGATG | Mutation sites detection of the *Aswhite* |
| *Aswhite* detection -w1-R | GGGAAACAGTAGCACCCAATA | |
| *Aswhite* detection-w2-F | GAACGATAAGGCGACGCTGAT | |
| *Aswhite* detection-w2-R | GGCGACGGGATGAACAGAT | |
| *Asyellow* detection-y1-F | CAGCCCGACTTCGTCTTCC | Mutation sites detection of the *Asyellow* |
| *Asyellow* detection-y1-R | TCGTGATAGGTGTCTTCTTCTCCA | |
| *Asyellow* detection-y2-F | ATAAGTGCGGACGCCTGTG | |
| *Asyellow* detection-y2-R | ATGCCGAAGATGCCCTCC | |
| P2C-Linker-Cas9-fragment1-F | GGTGCCGCGCGGCAGCCATATGCCAAAGAAGAAGCGGAAGGTCA | Construction of the *pET-28a-P2C-* |

| | | |
|---|---|---|
| P2C-Linker-Cas9-fragment1-R | TCTTGTCCATGCTCCCGCCTCCACCGCT | *Linker-Cas9* expression vector |
| P2C-Linker-Cas9-fragment2-F | AGGCGGGAGCATGGACAAGAAGTACTCCATTGGGC | |
| P2C-Linker-Cas9-fragment2-F | GTGGTGGTGGTGGTGCTCGAGCACCTTCCTCTTCTTCTTGGGG | |
| Cas9-F | CGCGCGGCAGCCATATGATGGACAAGAAGTACTCCATTGGGC | Construction of the *pET28a-Cas9* expression vector |
| Cas9-R | GGTGGTGGTGCTCGAGCACCTTCCTCTTCTTCTTGGGG | |
| DsRed-F | CGCGCGGCAGCCATATGATGGTGCGCTCCTCCAAG | Construction of the *pET28a-DsRed* expression vector |
| DsRed-R | GGTGGTGGTGCTCGAGCAGGAACAGGTGGTGGCG | |
| P2C-Linker-DsRed-fragment1-F | GGTGCCGCGCGGCAGCCATATGCCAAAGAAGAAGCGGAAGGTCA | Construction of the *pET28a-P2C-DsRed* expression vector |
| P2C-Linker-DsRed-fragment1-R | AGCGCACCATGCTCCCGCCTCCACCGCT | |
| P2C-Linker-DsRed-fragment2-F | AGGCGGGAGCATGGTGCGCTCCTCCAAG | |
| P2C-Linker-DsRed-fragment2-R | GTGGTGGTGGTGGTGCTCGAGCAGGAACAGGTGGTGGCG | |
| q*AsVg*-F | CCAGCCGAAGGTCTACATCAAC | RT-qPCR for *AsVg* |
| q*AsVg*-R | CTATGGTGGTGCCGCAAAGA | |
| q*AsRPL49*-F | GGAGCCGGTCGGTGATATGT | |
| q*AsRPL49*-R | TTCCTTCTCGGTCGGCTTCG | |

Note: The underlined letters represent the T7 promoter sequence, and the italicized letters represents the gRNA target site sequence.

Table S2. Survival rate and red fluorescence rate statistics of *Anopheles sinensis* after injection of pET28a-P2C-DsRed, pET28a-DsRed fusion proteins

| | Number of injected individuals | Survival individuals after 20 h of injection | Survival rate after 20 h of injection | red fluorescence individuals | red fluorescence rate |
|---|---|---|---|---|---|
| pET28a-P2C-DsRed | 27 | 27 | 100% | 27 | 100% |
| | 30 | 30 | 100% | 30 | 100% |
| pET28a-DsRed | 33 | 33 | 100% | 0 | 0 |
| | 28 | 28 | 100% | 0 | 0 |

Table S3. The number of laid eggs and the hatching rate after injection of a mixture of different concentrations of saponin and P2C-Linker-Cas9 protein into female *Anopheles sinensis* (at 20 h PBF)

| Mixture types | Mixture concentrations | Number of injected individuals | Number of surviving individuals at 20 hours after of injection | Survival rate at 20 hours after of injection | Number of laid eggs | Number of hatched eggs | Hatching rate |
|---|---|---|---|---|---|---|---|
| P2C-Linker-Cas9+ Saponin | 100ng/μL-1 | 10 | 10 | 100% | 1125 | 936 | 83.2% |
| | 100ng/μL-2 | 10 | 10 | 100% | 835 | 701 | 83.9% |
| | 150ng/μL-1 | 10 | 10 | 100% | 892 | 674 | 75.6% |
| | 150ng/μL-2 | 10 | 10 | 100% | 965 | 736 | 76.2% |
| | 200ng/μL-1 | 10 | 10 | 100% | 1083 | 791 | 73% |
| | 200ng/μL-2 | 10 | 10 | 100% | 868 | 647 | 74.5% |
| | 250ng/μL-1 | 10 | 10 | 100% | 837 | 598 | 71.4% |
| | 250ng/μL-2 | 10 | 10 | 100% | 922 | 650 | 70.5% |
| | 300ng/μL-1 | 10 | 9 | 90% | 1098 | 661 | 60% |
| | 300ng/μL-2 | 10 | 10 | 100% | 989 | 631 | 63.8% |
| | 350ng/μL-1 | 10 | 10 | 100% | 486 | 214 | 44% |
| | 350ng/μL-2 | 10 | 10 | 100% | 483 | 242 | 50% |
| P2C-Linker-Cas9 | 2500ng/μL-1 | 10 | 10 | 100% | 1066 | 951 | 89.2% |
| | 2500ng/μL-2 | 10 | 10 | 100% | 1042 | 984 | 94.4% |

Table S4. $G_0$ Gene-editing efficiency (GEF) after knock out of *Aswhite* gene (RNP complex were injected into the WT females which mated with WT males, named as WF × WM form)

| | P | | | $G_0$ | | | | | |
|---|---|---|---|---|---|---|---|---|---|
| | Number of injected individuals | Number of survival individuals | Survival rate | Number of Offspring Individuals | Number of individuals with mosaic phenotype | Number of individuals with white-eye phenotype | $G_0$ Gene-editing efficiency (GEF) | GEF (for *white*) in *Aedes aegypti** | GEF (for *white*) in *Culex pipiens pallens*+ |
| *Aswhite*1 | 124 | 102 | 82.2% | 3509 | 451 | 17 | 13.3% | 0.1% | 0.4%[a] |
| *Aswhite*2 | 126 | 105 | 83.3% | 3198 | 421 | 13 | 13.6% | | |
| Control 1 | 28 | 26 | 92.8% | 617 | 7 | 0 | 1.1% | | |
| Control 2 | 24 | 23 | 95.8% | 664 | 6 | 0 | 0.9% | | |

Note: WF, wild-type female; WM, wild-type male. The '*' symbol indicates the reference from Chaverra-Rodriguez's study. The '+' indicates the reference from Li's study. 'a' indicates the highest efficiency in the cited study.

Table S5. Phenotypes of the mutant $G_1$ offspring, resulting from self-crossing of $G_0$ (under the form of WF × WM)

| | WF × WM | |
|---|---|---|
| | $G_0$ | $G_1$ |
| | mosaic white ♀ × mosaic white ♂ | white mutant (♂ and ♀) √ |

| | | |
|---|---|---|
| P2C-Cas9-gRNA Injection group | full white♀× full white♂ | white mutant (♂ and ♀) √ |
| | mosaic yellow ♀× full yellow ♂ | yellow mutant (♂ and ♀) √ |
| | full yellow ♀× full yellow ♂ | yellow mutant (♂ and ♀) √ |
| Cas9-gRNA Injection group | mosaic white ♂ × mosaic white ♀ | white mutant (♂ and ♀) × |

Note: WF, wild-type female; WM, wild-type male. The '√' symbol indicates the presence of corresponding mutant phenotype individuals in the $G_1$ offspring, while the '×' symbol indicates the absence of corresponding mutant phenotype individuals in the $G_1$ offspring.

Table S6. $G_0$ Gene-editing efficiency (GEF) after knock out of *Asyellow* gene (RNP complex were injected into the WT females which mated with WT males, named as WF × WM form)

| | P | | | $G_0$ | | | | | |
|---|---|---|---|---|---|---|---|---|---|
| | Number of injected individuals | Number of survival individuals | Survival rate | Number of Offspring Individuals | Number of individuals with mosaic phenotype | Number of individuals with yellow-body phenotype | $G_0$ Gene-editing efficiency (GEF) | GEF (for *white*) in *Aedes aegypti*[*] | GEF (for *white*) in *Culex pipiens pallens*[+] |
| *Asyellow*1 | 184 | 174 | 94.5% | 8324 | 27 | 29 | 0.7% | 0.1% | 0.4%[a] |
| *Asyellow*2 | 163 | 160 | 98.1% | 7643 | 18 | 19 | 0.5% | | |
| Control 1 | 56 | 56 | 100% | 3505 | 0 | 0 | 0 | | |
| Control 2 | 45 | 41 | 91% | 3212 | 0 | 0 | 0 | | |

Note: WF, wild-type female; WM, wild-type male. The '*' symbol indicates the reference from Chaverra-Rodriguez's study. The '+' indicates the reference from Li's study. 'a' indicates the highest efficiency in the cited study.

Table S7. $G_0$ Gene-editing efficiency (GEF) after knock out of *Aswhite* gene (RNP complex were injected into the WT females which mated with mutant WT males, named as WF × MM form)

| | P | | | $G_0$ | | | | | |
|---|---|---|---|---|---|---|---|---|---|
| | Number of injected individuals | Number of survival individuals | Survival rate | Number of Offspring Individuals | Number of individuals with mosaic phenotype | Number of individuals with white - eye phenotype | $G_0$ Gene-editing efficiency (GEF) | GEF (for *white*) in *Aedes aegypti*[*] | GEF (for *ECFP* in transgenic line) in *Anopheles stephensi*[+#] |
| *Aswhite*1 | 180 | 141 | 78.3% | 5763 | 1827 | 97 | 33.4% | 2.4%[a] | 5.6%[a] |
| *Aswhite*2 | 173 | 144 | 83.2% | 5997 | 1709 | 77 | 29.8% | | |
| Control 1 | 47 | 44 | 93.6% | 2502 | 10 | 0 | 0.4% | | |
| Control 2 | 57 | 51 | 89.4% | 2712 | 14 | 0 | 0.5% | | |

Note: WF, wild-type female; MM, mutant male. The '*' symbol indicates the reference from Chaverra-Rodriguez's study. The ‘+’ indicates the reference from Macias’s study. [#] It should be noted that the researchers used ECFP transgenic females to mate with wild-type males and injected the Cas9-ECFP gRNA complex into the females. The editing efficiency was analyzed by counting the proportion of offspring that did not exhibit ECFP fluorescence. This approach is consistent with our study and belongs to the type of material gene editing. ‘a’ indicates the highest efficiency in the cited study.

Table S8. $G_0$ Gene-editing efficiency (GEF) after knock out of *Asyellow* gene (RNP complex were injected into the WT females which mated with mutant males, named as WF × MM form)

| | P | | | | $G_0$ | | | | |
|---|---|---|---|---|---|---|---|---|---|
| | Number of injected individuals | Number of survival individuals | Survival rate | Number of Offspring Individuals | Number of individuals with mosaic phenotype | Number of individuals with yellow-body phenotype | $G_0$ Gene-editing efficiency (GEF) | GEF (for *white*) in *Aedes aegypti*[*] | GEF (for *ECFP* in transgenic line) in *Anopheles stephensi*[+#] |
| *Asyellow*1 | 47 | 42 | 89% | 2562 | 30 | 69 | 3.9% | 2.4%[a] | 5.6%[a] |
| *Asyellow*2 | 77 | 71 | 92.2% | 3763 | 34 | 93 | 34% | | |
| Control 1 | 58 | 56 | 96.5% | 3412 | 0 | 0 | 0 | | |
| Control 2 | 49 | 44 | 89.8% | 3063 | 0 | 0 | 0 | | |

Note: WF, wild-type female; MM, mutant male. The ‘*’ symbol indicates the reference from Chaverra-Rodriguez’s study. The ‘+’ indicates the reference from Macias’s study. [#] It should be noted that the researchers used ECFP transgenic females to mate with wild-type males and injected the Cas9-ECFP gRNA complex into the females. The editing efficiency was analyzed by counting the proportion of offspring that did not exhibit ECFP fluorescence. This approach is consistent with our study and belongs to the type of material gene editing. ‘a’ indicates the highest efficiency in the cited study.

Table S9. Phenotypes of the mutant $G_1$ offspring, resulting from self-crossing of $G_0$ (under the form of WF × MM)

| WF × MM | | |
|---|---|---|
| | $G_0$ | $G_1$ |
| P2C-Cas9-gRNA Injection group | mosaic white ♀ × mosaic white♂ | white mutant (♂ and ♀) √ |
| | full white ♀ × full white♂ | white mutant (♂ and ♀) √ |
| | mosaic yellow ♀ × mosaic yellow♂ | yellow mutant (♂ and ♀) √ |
| | full yellow ♀ × full yellow♂ | yellow mutant (♂ and ♀) √ |
| Cas9-gRNA Injection group | mosaic white ♀ ×mosaic white♂ | white mutant (♂ and ♀) × |

Note: WF, wild-type female; WM, wild-type male. The ‘√’ symbol indicates the presence of corresponding mutant phenotype individuals in the $G_1$ offspring, while the ‘×’ symbol indicates the absence of corresponding mutant phenotype individuals in the $G_1$ offspring.
