## Supplementary figures and images for "High-efficiency gene editing in *Anopheles sinensis* using ReMOT control"

### 20230821 Fig S1.pdf

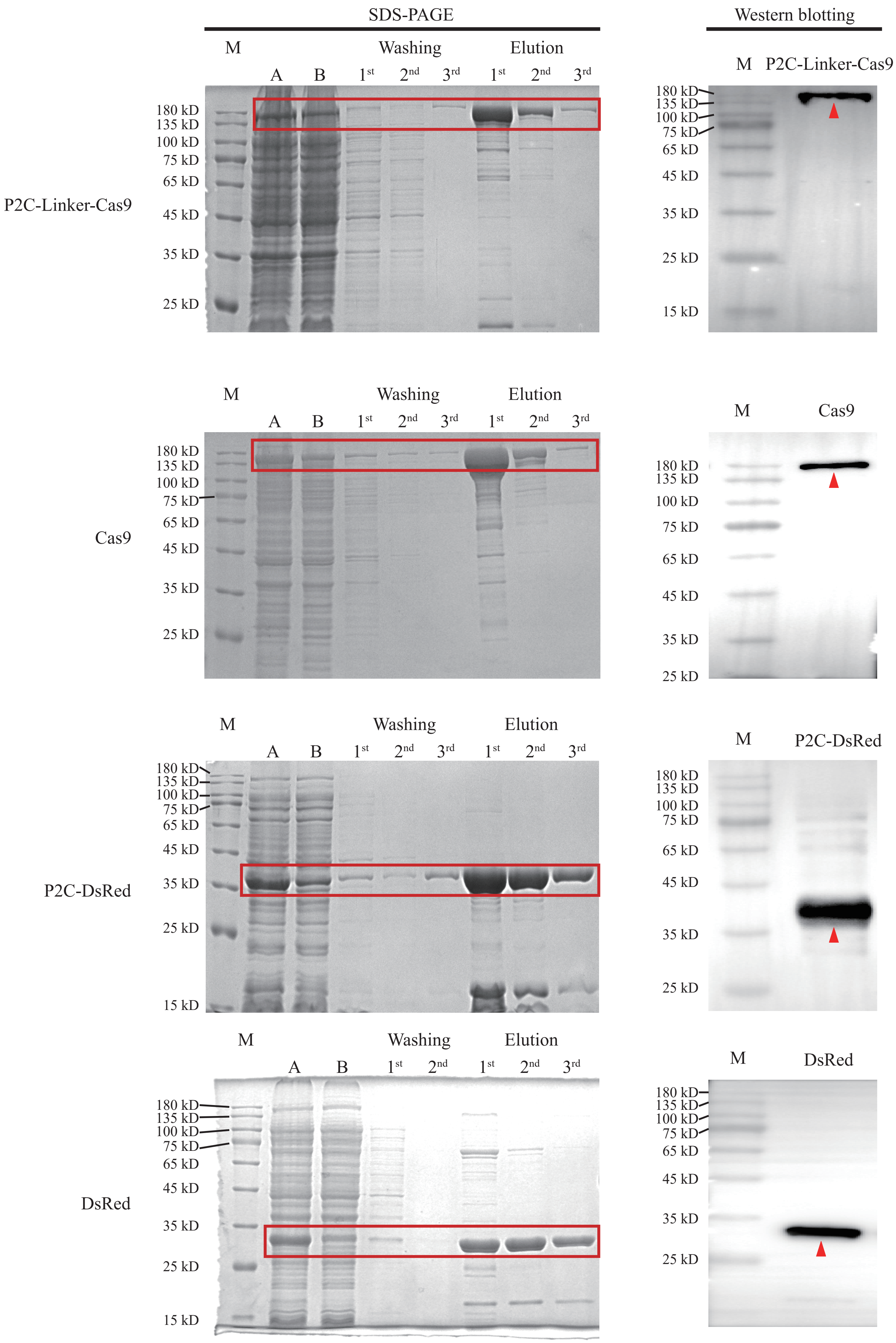

### 20230821 Fig S2.pdf

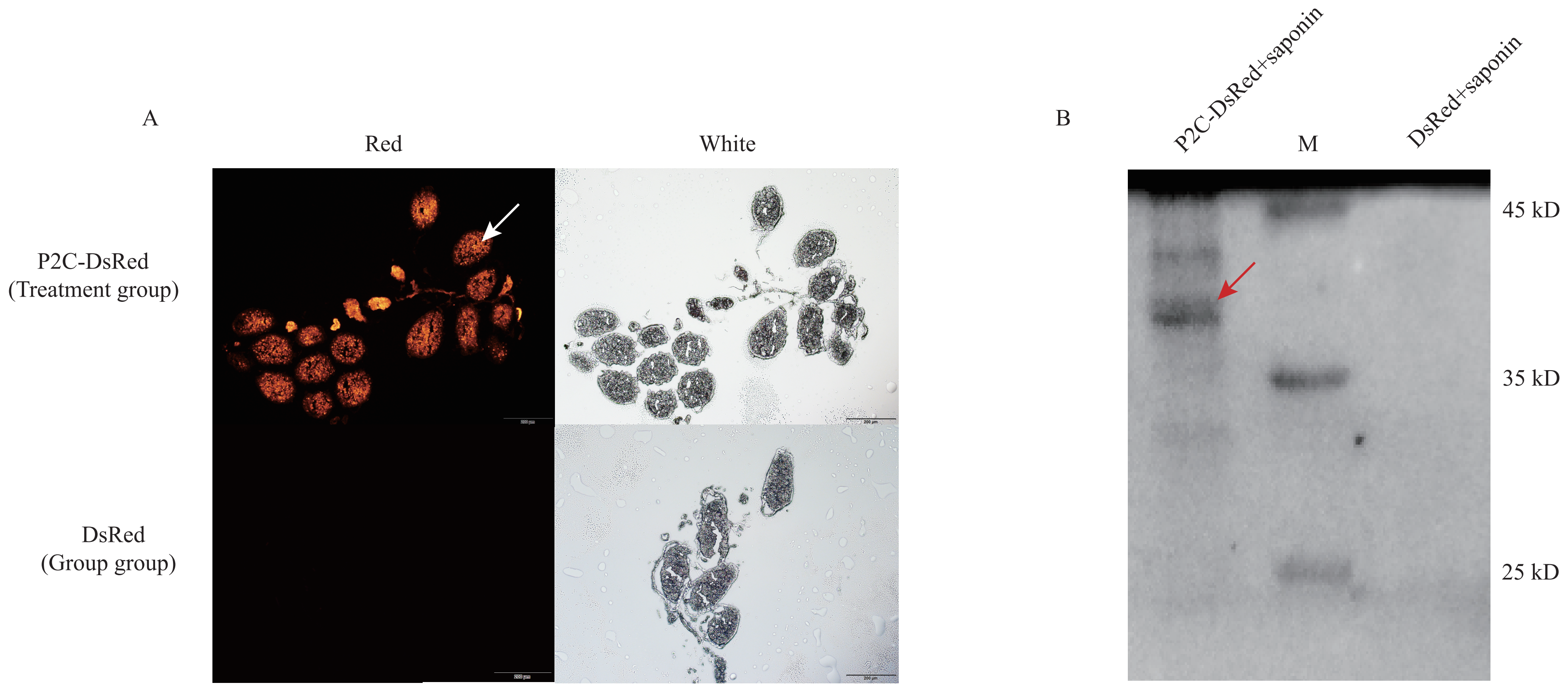

### 20230821 Fig S3.pdf

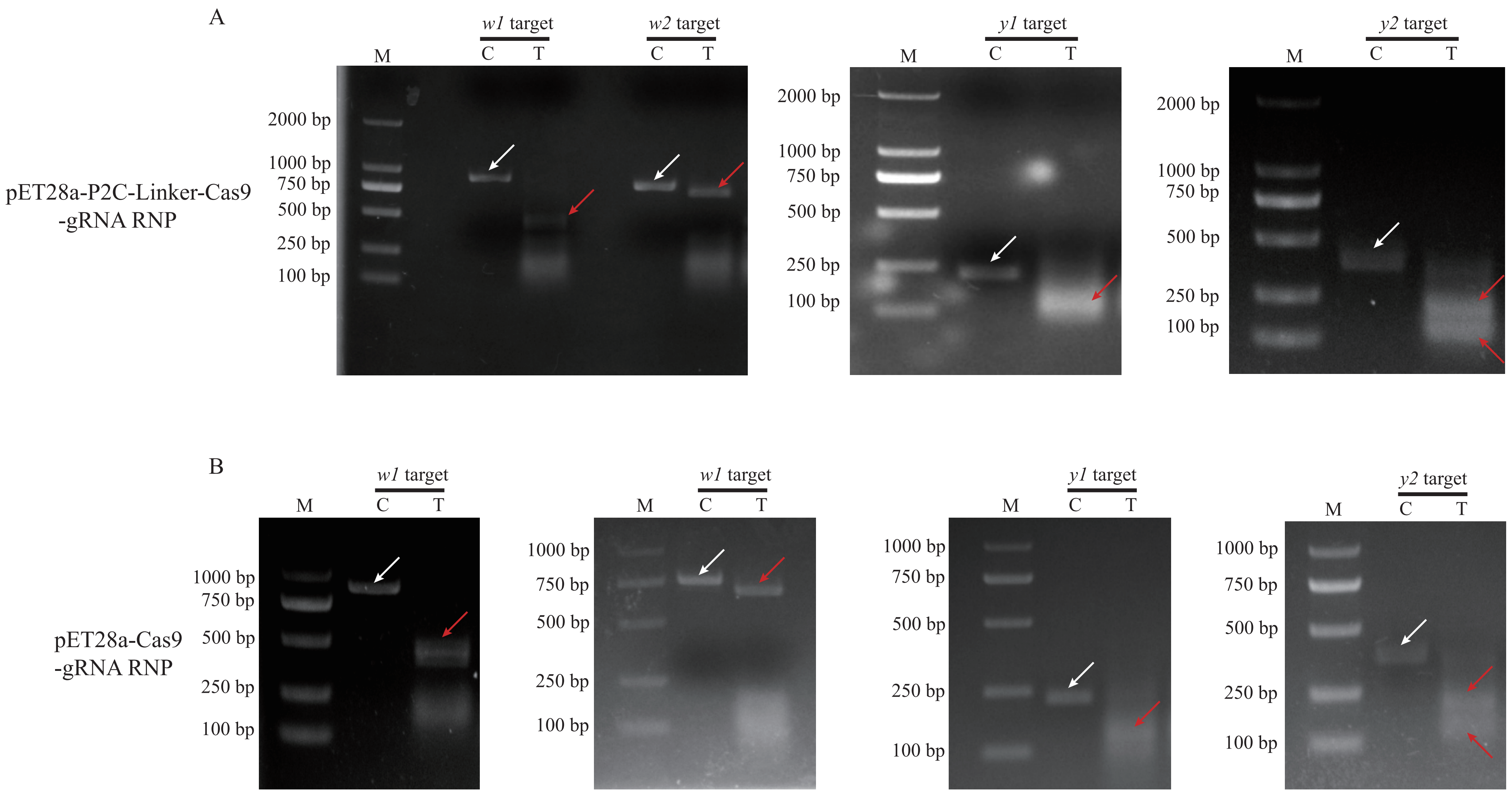
